# Experimental and natural evidence of SARS-CoV-2 infection-induced activation of type I interferon responses

**DOI:** 10.1101/2020.06.18.158154

**Authors:** Arinjay Banerjee, Nader El-Sayes, Patrick Budylowski, Daniel Richard, Hassaan Maan, Jennifer A. Aguiar, Kaushal Baid, Michael R. D’Agostino, Jann Catherine Ang, Benjamin J.-M. Tremblay, Sam Afkhami, Mehran Karimzadeh, Aaron T. Irving, Lily Yip, Mario Ostrowski, Jeremy A. Hirota, Robert Kozak, Terence D. Capellini, Matthew S. Miller, Bo Wang, Samira Mubareka, Allison J. McGeer, Andrew G. McArthur, Andrew C. Doxey, Karen Mossman

**Author notes:** These authors contributed equally (Nader El-Sayes, Patrick Budylowski). Correspondence (A.B.), (K.M.). 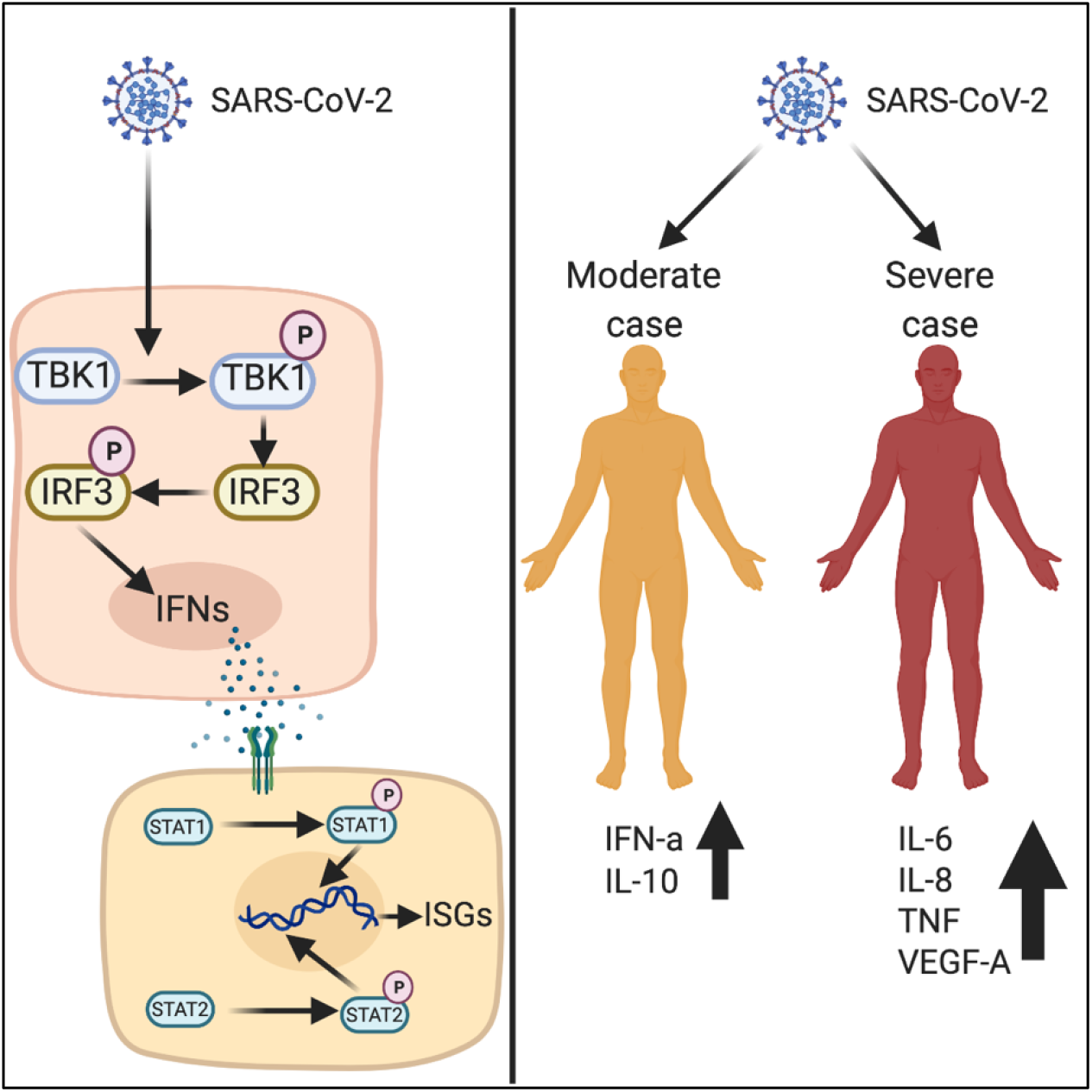 GRAPHICAL SUMMARY. **HIGHLIGHTS** - SARS-CoV-2 infection in human lung cells induces the expression of type I interferons - SARS-CoV-2 infection activates canonical transcription factors that are involved in type I interferon expression and signaling - SARS-CoV-2 cannot inhibit exogenous stimulation of type I IFN expression - SARS-CoV-2 cannot inhibit exogenous activation of type I IFN signaling - Moderate cases of COVID-19 upregulate higher serum levels of IL10 and IFNα, whereas severe cases of COVID-19 display higher serum levels of IL6, TNFα and IL8.

## Abstract

Type I interferons (IFNs) are our first line of defence against a virus. Protein over-expression studies have suggested the ability of SARS-CoV-2 proteins to block IFN responses. Emerging data also suggest that timing and extent of IFN production is associated with manifestation of COVID-19 severity. In spite of progress in understanding how SARS-CoV-2 activates antiviral responses, mechanistic studies into wildtype SARS-CoV-2-mediated induction and inhibition of human type I IFN responses are lacking. Here we demonstrate that SARS-CoV-2 infection induces a mild type I IFN response *in vitro* and in moderate cases of COVID-19. *In vitro* stimulation of type I IFN expression and signaling in human airway epithelial cells is associated with activation of canonical transcriptions factors, and SARS-CoV-2 is unable to inhibit exogenous induction of these responses. Our data demonstrate that SARS-CoV-2 is not adept in blocking type I IFN responses and provide support for ongoing IFN clinical trials.

## Introduction

Severe acute respiratory syndrome coronavirus 2 (SARS-CoV-2) emerged in December 2019 to cause a global pandemic of coronavirus disease (COVID-19) (Zhou et al., 2020a). SARS-CoV-2 causes a respiratory infection, along with acute respiratory distress syndrome in severe cases. Innate antiviral responses, which include type I interferons (IFNs), are the first line of antiviral defense against an invading virus (Kawai and Akira, 2006). Cellular pattern recognition receptors (PRRs) recognize viral nucleic acids and activate key cellular kinases, such as Inhibitor of nuclear factor Kappa-B Kinase subunit epsilon (IKKε) and TANK-Binding Kinase 1 (TBK1).

These kinases phosphorylate and activate transcription factors such as interferon regulatory factor 3 (IRF3) to stimulate downstream production of type I/III IFNs (Koyama et al., 2008). Type I IFNs interact with interferon alpha/beta receptor (IFNAR) on cells to induce phosphorylation and activation of downstream mediators, such as signal transducer and activator of transcription 1 and 2 (STAT1 and STAT2), which leads to the production of antiviral interferon stimulated genes (ISGs). Similarly, Type III IFNs interact with their cognate receptors, IL-10R2 and IFNLR1 to activate STAT1 and STAT2, followed by the production of ISGs (Mesev et al., 2019).

Viruses encode proteins that can inhibit type I IFN production and signaling (Katze et al., 2002; Schulz and Mossman, 2016). Emerging pathogenic human coronaviruses, such as SARS-CoV and Middle East respiratory syndrome (MERS)-CoV have evolved multiple proteins that inhibit type I IFN responses in human cells (Chen et al., 2014; de Wit et al., 2016; Lu et al., 2011; Siu et al., 2014; Yang et al., 2013). Thus, to better understand SARS-CoV-2 pathogenesis, it is critical to identify the dynamic interaction of SARS-CoV-2 and the type I IFN response. Emerging data suggest that ectopic expression of at least 13 SARS-CoV-2 proteins, namely NSP1, NSP3, NSP6, NSP12, NSP13, NSP14, NSP15, M, ORF3a, ORF6, ORF7a, ORF7b and ORF9b can inhibit type I IFN responses in human cells (Gordon et al., 2020; Jiang et al., 2020; Lei et al., 2020; Xia et al., 2020). However, these data were largely derived from over-expression studies where SARS-CoV-2 proteins were selectively over-expressed in human cells to identify their immune response modulation capabilities. Studies based on over-expression of viral proteins do not mirror the dynamics of viral gene expression and their timing- and dose-dependent effects on host signaling. These dynamic events include activation of type I IFN responses via generation of viral pathogen associated molecular patterns (PAMPs), such as double-stranded RNA (dsRNA), followed by subsequent modulation of this antiviral response by viral proteins. Indeed, contradictory observations have been reported by studies on SARS-CoV-2 proteins that have employed an over-expression model to identify host protein interacting partners. For example, SARS-CoV-2 NSP15 has been reported as an IFN-modulating protein by Gordon *et al.* (Gordon et al., 2020), but Lei *et al.* (Lei et al., 2020) were unable to identify NSP15 as an inhibitor of IFN promoter activation. In addition, both Gordon *et al.* and Jiang *et al.* identified ORF9b as a modulator of IFN responses (Gordon et al., 2020; Jiang et al., 2020; Lei et al., 2020), but the study by Lei *et al.* did not identify ORF9b as a modulator (Lei et al., 2020). Furthermore, infection with wildtype SARS-CoV-2 in Caco-2 cells activated phosphorylation of TBK1 and IRF3, along with mild induction of ISGs (Shin et al., 2020). Thus, in-depth studies with clinical isolates of SARS-CoV-2 are required to confidently identify type I IFN responses that are generated in infected human cells and to determine if infection with wildtype virus isolates can sufficiently counteract these protective antiviral responses.

Transcriptional data from *in vitro* and *in vivo* work have demonstrated the lack of induction of type I IFN responses following SARS-CoV-2 infection (Blanco-Melo et al., 2020). In contrast, emerging data from patients with mild and moderate cases of COVID-19 have demonstrated the presence of type I IFN (Hadjadj et al., 2020a; Trouillet-Assant et al., 2020). Subsequently, recent studies have identified robust type I IFN responses in severe COVID-19 cases, which have been speculated to be associated with an exacerbated inflammatory response (Zhou et al., 2020b). In addition, upregulation of ISGs was also identified in a single-cell RNA sequencing study of peripheral blood mononuclear cells (PBMCs) from hospitalized COVID-19 patients (Wilk et al., 2020). Studies with patient samples are critical to understand the pathogenesis of SARS-CoV-2; however, the timing of sample collection, case definition of disease severity and varying viral load can lead to different observations related to IFN responses. An early and controlled IFN response is preferable during virus infection. Excessive induction of type I IFN responses in COVID-19 patients is associated with higher levels of damaging inflammatory molecules (Lucas et al., 2020). Thus, it is critical to identify the extent to which SARS-CoV-2 can induce or inhibit human IFN responses using controlled and robust mechanistic studies.

In this study, we have identified global early transcriptional responses that are initiated during infection of human lung epithelial (Calu-3) cells at 0, 1, 2, 3, 6, and 12 hours post incubation with a clinical isolate of SARS-CoV-2 from a COVID-19 patient in Toronto (Banerjee et al., 2020a). Data from our study demonstrate that *in vitro* SARS-CoV-2 infection induces the expression of type I IFNs, along with the expression of downstream ISGs. We also identified an increasing trend for type I IFN expression (IFN-α2) in sera from moderate cases of COVID-19, relative to healthy individuals and severe cases of COVID-19. We performed mechanistic studies to identify the mode of activation of type I IFNs using a clinical isolate of SARS-CoV-2. *In vitro* infection with SARS-CoV-2 induced phosphorylation of canonical transcription factors that are involved in the type I IFN response, such as IRF3, STAT1 and STAT2; exogenous activation of these transcription factors was not inhibited by wildtype SARS-CoV-2. In addition, we detected higher serum levels of anti-inflammatory cytokines in moderate cases of COVID-19 than in severe cases. Severe cases of COVID-19 displayed higher serum levels of pro-inflammatory cytokines. Data from our study demonstrate that wildtype SARS-CoV-2 is unable to block IFN responses. Further mechanistic studies are warranted to identify host factors that contribute to varying disease severity during the course of COVID-19, along with the regulation of inflammatory and anti-inflammatory cellular processes in SARS-CoV-2 infected cells.

## Results

### Global cellular response in SARS-CoV-2 infected human airway epithelial cells

The replication cycle of CoVs is complex and involves the generation of sub-genomic RNA molecules, which in turn code for mRNA that are translated into proteins (Banerjee et al., 2019; Sawicki et al., 2007). To determine SARS-CoV-2 replication kinetics in human cells using RNA-seq, we infected human airway epithelial cells (Calu-3) at a multiplicity of infection (MOI) of 2. After incubation with virus inoculum for 1-hour, media was replaced with cell growth media and RNA was extracted and sequenced (poly-A enriched RNA) at 0-, 1-, 2-, 3-, 6- and 12-hours post incubation (hpi). SARS-CoV-2 genome, sub-genomic RNA and transcripts were detected in infected samples; viral transcript expression clustered based on post-incubation time using principal component analysis (PCA) (see supplementary Figure S1A). In our RNA-seq analysis, we detected high levels of expression of SARS-CoV-2 structural and accessory genes at the 3’ end of the genome as early as 0 hpi (Figure 1A). Significant expression of *ORF1ab*, relative to 0 hpi was detected at 6 hpi (Figure 1B). SARS-CoV-2 nucleocapsid (*N*) gene was highly expressed relative to other genes as early as 0 hpi (Figure 1B), with relative expression significantly increasing over time (*p*=1.4e-16; Figure 1B). The absolute expression of other genes increased over time with levels of *N* > *M* > *ORF10* > *S* > *ORF1ab* > *ORF7a* > *ORF8* > *ORF3a* > *ORF6* > *E* > *ORF7b* > *ORF1a* at 12 hpi (Figure 1B and supplementary Table S1).

**Figure 1.**
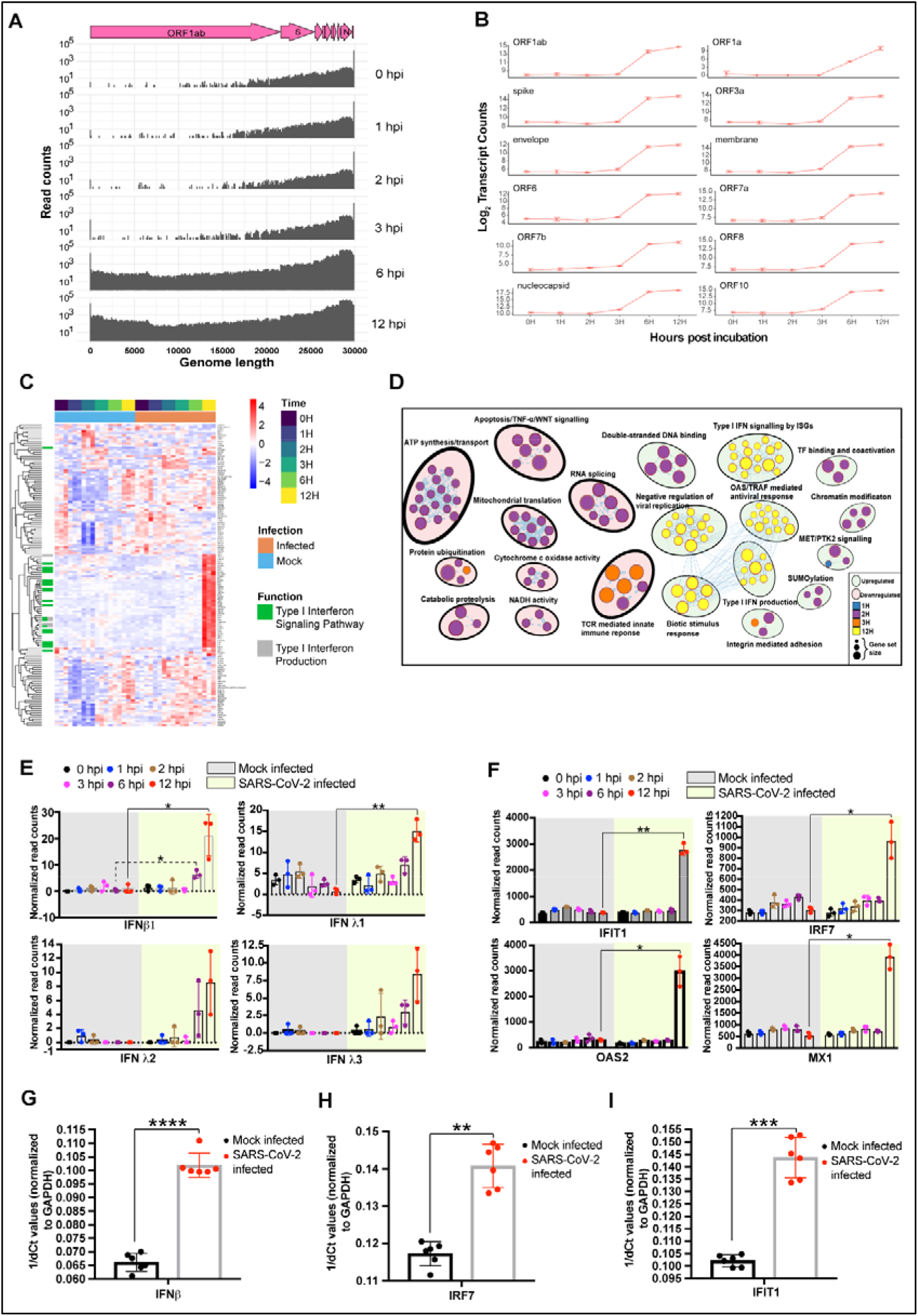
Global response in SARS-CoV-2 infected human airway epithelial cells. Calu-3 cells were infected with SARS-CoV-2 at an MOI of 1 or 2. RNA was extracted at different times post incubation. Viral and cellular gene expression was determined using time-series RNA-seq analysis or qPCR. (**A**) SARS-CoV-2 gene expression over 12 hours (n = 3/time point). The genome organization of SARS-CoV-2 is indicated above in pink. (**B**) Major SARS-CoV-2 gene expression levels at different times post incubation (n = 3/time point). Statistical analysis was performed in R (see methods). (**C**) Cellular genes (n = 124) that are significantly up or downregulated (FDR-adjusted *p*<0.05; |log_2_FC| > 1) in SARS-CoV-2 infected cells, relative to mock infected cells at different times post incubation. Transcript levels are shown as z-score normalized expression (scaled by gene). See supplementary Figure S1E for a larger figure. (**D**) Cellular processes that are down or upregulated at different times post incubation. The size of the circles represents the number of genes that are down or upregulated at different times after incubation (n = 3/time point). (**E**) Transcript abundance of type I interferon (IFN) genes (*IFNβ* and *IFNλ1-3*) in mock infected and SARS-CoV-2 infected Calu-3 cells at different times (n = 3). (**F**) Transcript abundance of representative interferon stimulated genes (ISGs) in mock infected and SARS-CoV-2 infected Calu-3 cells at different times (n = 3). (**G**) *IFNβ* transcript levels in Calu-3 cells infected with SARS-CoV-2 or mock infected for 12 hours, normalized to *GAPDH* (n = 6). Transcript levels were determined by qPCR. (**H**) *IRF7* transcript levels in Calu-3 cells infected with SARS-CoV-2 or mock infected for 12 hours, normalized to *GAPDH* (n = 6). Transcript levels were determined by qPCR. (**I**) *IFIT1* transcript levels in Calu-3 cells infected with SARS-CoV-2 or mock infected for 12 hours, normalized to *GAPDH* (n = 6). Transcript levels were determined by qPCR. Data are represented as mean ± SD, n = 3 or 6, *p*\*<0.05, **<0.01, ***<0.001 and ****<0.0001 (Student’s t test). See also Transparent Methods for details on statistical analyses performed using *R*. See also supplementary Figures S1–S3, and supplementary Tables S1–S3. H and hpi, hours post incubation. See also supplementary Figures S1 and S2, and supplementary Tables S1–3.

To determine SARS-CoV-2 infection-mediated host responses, we extracted total cellular RNA at different times post infection and analyzed gene expression in infected and mock infected Calu-3 cells using RNA-seq. Gene expression levels in these cells clustered based on time-points via PCA (see supplementary Figure S1B). One hundred and twenty-four genes were significantly differentially expressed in infected cells (FDR-adjusted *p* < 0.05), relative to mock infected cells in at least one time point post infection (|log_2_FC| > 1), including genes involved in type I IFN production and signaling (Figure 1C; see supplementary Table S2 and Figures S1C and S1E). The extent of antiviral gene expression at 12 hpi correlated with an increase in viral transcripts (see supplementary Figure S1C). Interestingly, at early time points of 2 and 3 hpi, pathway enrichment analysis revealed numerous cellular processes that were significantly downregulated in SARS-CoV-2 infected cells, relative to mock infected cells (FDR-adjusted *p*<0.05). Downregulated processes included RNA splicing, apoptosis, ATP synthesis and host translation, while genes associated with viral processes, cell adhesion and double-stranded RNA binding were upregulated in infected cells relative to mock infected cells at 2 and 3 hpi (Figure 1D; see supplementary Figures S1D and S2, and supplementary Table S3). Cellular pathways associated with type I IFN production and signaling, along with OAS/TRAF-mediated antiviral responses were significantly upregulated at 12 hpi (Figure 1D and see supplementary Figure S2). Consistent with other reports (Blanco-Melo et al., 2020), transcript levels for *IFNβ1* and *IFNλ1* were significantly upregulated at 12 hpi with SARS-CoV-2 (Figure 1E). Transcript levels of *IFNλ2* and *IFNλ3* were elevated at 6 and 12 hpi, but the levels did not reach significance relative to mock infected cells at these time points (Figure 1E).

IFN production alone is not sufficient to protect cells from invading viruses. IFNs function through ISG expression, which in turn confers antiviral protection in infected (autocrine mode of action) and neighbouring (paracrine mode of action) cells (Schoggins, 2019; Schoggins and Rice, 2011). Nineteen antiviral ISGs were upregulated in infected cells, relative to mock infected cells at 12 hpi, including interferon induced protein with tetratricopeptide repeats 1 (*IFIT1*), interferon regulatory factor 7 (*IRF7*), 2’-5-oligoadenylate synthetase 2 (*OAS2*) and MX dynamin GTPase 1 (*MX1*) (Figure 1F; see supplementary Figure S3A and supplementary Table S2). Genes associated with structural molecule activity, cell adhesion and exocytosis were downregulated in SARS-CoV-2 infected cells, relative to uninfected cells at 12 hpi (see supplementary Figure S2).

Coronaviruses, such as those that cause SARS and MERS have evolved multiple proteins that can inhibit type I IFN expression (Chen et al., 2014; Lu et al., 2011; Lui et al., 2016; Niemeyer et al., 2013; Siu et al., 2014; Yang et al., 2013). To confirm our RNA-seq findings that SARS-CoV-2 infection alone is sufficient to induce type I IFN and ISG responses in Calu-3 cells, we infected cells with SARS-CoV-2 and assessed transcript levels of *IFNβ*, *IRF7* and *IFIT1* by quantitative polymerase chain reaction (qPCR). *IFNβ* induction was observed 12 hpi in SARS-CoV-2 infected cells, relative to mock-infected cells (Figure 1G). Consistent with the upregulation of *IFNβ* transcripts in SARS-CoV-2 infected cells, transcript levels for ISGs, such as *IRF7* and *IFIT1* were also significantly upregulated at 12 hpi relative to mock infected cells (Figures 1H and 1I).

### SARS-CoV-2 infection fails to inhibit exogenous stimulation of type I IFN expression

To determine if SARS-CoV-2 is able to inhibit type I IFN responses mounted against an exogenous stimulus, we infected Calu-3 cells with SARS-CoV-2 for 12 hours at a MOI of 1 and stimulated these cells with exogenous double-stranded RNA [poly(I:C)] for 6 hours. We quantified SARS-CoV-2 replication by qPCR using primers designed to amplify genomic RNA by targeting a region between *ORF3a* and *E* genes. We called this region ‘upstream of E’ (UpE). SARS-CoV-2 UpE levels were higher in SARS-CoV-2 infected cells and in SARS-CoV-2 infected + poly(I:C) treated cells, relative to UpE levels at 0 hpi immediately after removing the inoculum (Figure 2A). We also measured the levels of *IFNβ* transcripts in these cells by qPCR. Poly(I:C) transfection alone induced higher levels of *IFNβ* transcripts relative to mock transfected cells (Figure 2B). SARS-CoV-2 infection alone also induced higher levels of *IFNβ* transcripts relative to mock infected cells (Figure 2B). Interestingly, there was no significant difference in *IFNβ* transcript levels between poly(I:C) transfected and SARS-CoV-2 infected + poly(I:C) transfected cells (Figure 2B).

**Figure 2.**
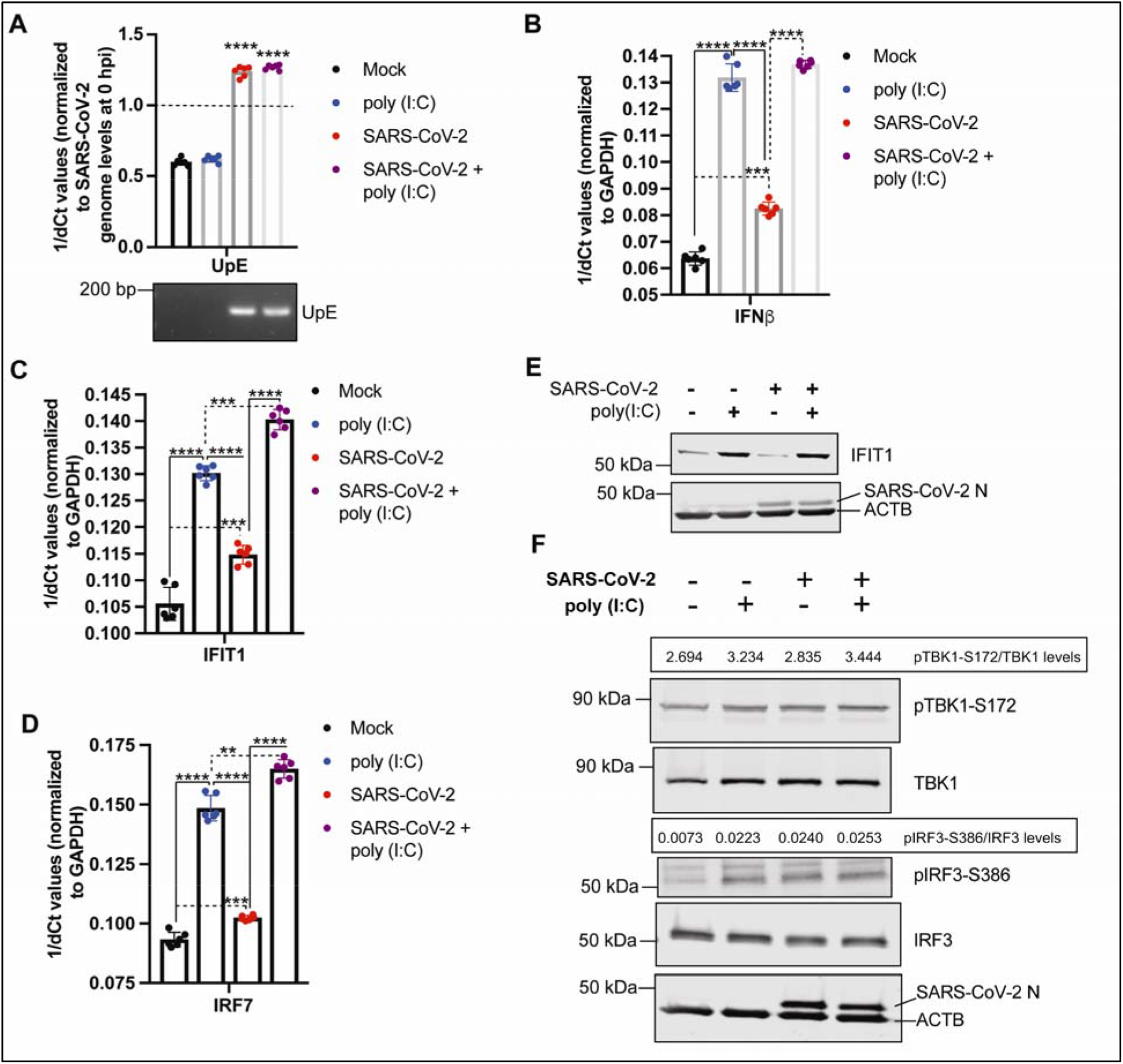
SARS-CoV-2 infection cannot inhibit type I IFN expression. To determine if SARS-CoV-2 can modulate *IFNβ* gene expression and downstream stimulation of ISGs, Calu-3 cells were infected with SARS-CoV-2 at an MOI of 1 for varying times, following which cells were mock transfected or transfected with poly(I:C). Mock infected and mock transfected cells served as controls. Transcript levels were quantified using qPCR. Protein expression was observed and quantified using immunoblots. (**A**) SARS-CoV-2 genome (UpE) levels in Calu-3 cells infected with SARS-CoV-2 or mock infected for 12 hours, and transfected with 100 ng of poly(I:C) or mock transfected for 6 hours (n = 6). Primers for the UpE region were designed to quantify SARS-CoV-2 genome levels (see Methods). 1/dCT values are represented after normalizing Ct values for SARS-CoV-2 genome levels at 18 hpi with Ct values observed at 0 hpi (immediately after removal of virus inoculum). Gel (below): UpE qPCR amplicons were also visualized on an agarose gel. (**B**) *IFNβ* transcript levels in Calu-3 cells that were infected with SARS-CoV-2 or mock infected for 12 hours. Twelve hpi, cells were either transfected with 100 ng of poly(I:C) or mock transfected for 6 hours. *IFNβ* transcript levels were normalized to *GAPDH* transcript levels (n = 6). (**C**) *IFIT1* transcript levels in Calu-3 cells that were infected with SARS-CoV-2 or mock infected for 12 hours. Twelve hpi, cells were either transfected with 100 ng of poly(I:C) or mock transfected for 6 hours. *IFIT1* transcript levels were normalized to *GAPDH* transcript levels (n = 6). (**D**) *IRF7* transcript levels in Calu-3 cells that were infected with SARS-CoV-2 or mock infected for 12 hours. Twelve hpi, cells were either transfected with 100 ng of poly(I:C) or mock transfected for 6 hours. *IRF7* transcript levels were normalized to *GAPDH* transcript levels (n = 6). (**E**) IFIT1, SARS-CoV-2 N and ACTB protein expression in Calu-3 cells that were infected with SARS-CoV-2 or mock infected for 24 hours. Twenty-four hpi, cells were either transfected with 1000 ng of poly(I:C) or mock transfected for 24 hours (n = 3). (**F**) pTBK1-S172, TBK1, pIRF3-S386, IRF3, SARS-CoV-2 N and ACTB protein expression in Calu-3 cells that were infected with SARS-CoV-2 or mock infected for 24 hours. Twenty-four hpi, cells were either transfected with 1000 ng of poly(I:C) or mock transfected for an additional 24 hours (n = 3). Data are represented as mean ± SD, n = 3 or 6, *p*\*\*<0.01, ***<0.001 and ****<0.0001 (Student’s t test). pTBK1-S172 and pIRF3-S386 protein expression levels are expressed as ratios of pTBK1-S172/TBK1 and pIRF3-S386/IRF3 levels, respectively. Blots were quantified using Image Studio (Li-COR) (n = 3). Ct, cycle threshold. See also supplementary Figure S3.

To determine if *IFNβ* expression in SARS-CoV-2 infected and/or poly(I:C) transfected cells is associated with ISG expression, we additionally quantified the levels of *IFIT1* and *IRF7*. Poly(I:C) transfection alone induced significantly higher levels of *IFIT1* and *IRF7* transcripts relative to mock transfected cells (Figures 2C and 2D). SARS-CoV-2 infection alone also induced higher levels of *IFIT1* and *IRF7* transcripts relative to mock infected cells (Figures 2C and 2D). Notably, *IFIT1* and *IRF7* transcript levels in SARS-CoV-2 infected + poly(I:C) transfected cells were higher than levels in cells that were transfected with poly(I:C) alone (Figures 2C and 2D), suggesting a lack of impact by SARS-CoV-2 on poly(I:C)-mediated induction.

For comparative purposes, we evaluated another respiratory RNA virus, influenza A virus (H1N1), which is known to both activate and modulate IFN responses. On comparison with H1N1 infection in Calu-3 cells, upregulation of *IFNβ*, *IFIT1* and *IRF7* transcript levels in SARS-CoV-2 infected cells seemed modest (Figure 2B-D and see supplementary Figure S3B). H1N1 infection with a similar MOI of 1 induced higher levels of *IFNβ* transcripts in Calu-3 cells relative to mock infected cells (see supplementary Figure S3B). H1N1 infection also induced higher levels of *IFIT1* and *IRF7* in Calu-3 cells relative to levels in mock infected cells (see supplementary Figure S3B). Importantly, H1N1 infection-induced upregulation of *IFIT1* and *IRF7* were comparable or greater than upregulation observed in poly(I:C) transfected cells (see supplementary Figure S3B), unlike in SARS-CoV-2 infected cells where *IFIT1* and *IRF7* expression was significantly lower than poly(I:C) transfected cells (Figures 2C and 2D). Infection with either SARS-CoV-2 or H1N1 was unable to inhibit poly(I:C)-mediated upregulation of *IFNβ*, *IFIT1* and *IRF7* transcript levels (Figures 2B-D and see supplementary Figure S3B).

To validate our gene expression observations, we examined SARS-CoV-2 N, IFIT1, and beta-actin (ACTB) protein expression. Poly(I:C) transfection induced higher levels of IFIT1 in Calu-3 cells, while SARS-CoV-2 infection did not induce higher observable levels of IFIT1 by immunoblot analysis at 48 hpi, relative to mock infected cells (Figure 2E); however, at 72 hpi, SARS-CoV-2 infection induced higher observable levels of IFIT1 protein expression relative to mock infected cells (see supplementary Figure S4). Consistent with our qPCR results, H1N1 infection alone induced detectable and higher levels of IFIT1 at 18 hpi (see supplementary Figure S3D). We confirmed SARS-CoV-2 and H1N1 infection in these cells by detecting N or NP protein in the samples, respectively (Figure 2E and supplementary Figure S3D). Both SARS-CoV-2 and H1N1 failed to inhibit the expression of poly(I:C)-induced IFIT1 (Figure 2E and supplementary Figure S3D).

Type I IFN production is primarily mediated by the phosphorylation and activation of TBK1, which in turn phosphorylates and activates IRF3 (Janeway and Medzhitov, 2002; Kawai and Akira, 2006). Activation of TBK1 is associated with phosphorylation of serine 172 (Larabi et al., 2013), while activation of IRF3 involves phosphorylation of serine 386, amongst other residues (Chen et al., 2008). To determine SARS-CoV-2 infection-induced phosphorylation of TBK1 and IRF3, we infected Calu-3 cells for 24 hours followed by poly(I:C) or mock stimulation for a further 24 hours and performed immunoblot analysis to detect levels of TBK1 (pTBK1-S172) and IRF3 (pIRF3-S386) phosphorylation. Only modest increases in phosphorylation of TBK1 were observed in SARS-CoV-2 infected and poly(I:C) treated cells relative to untreated cells. The highest level of pTBK1-S172 was observed in SARS-CoV-2 infected + poly(I:C) cells, followed by poly(I:C) treated cells (Figure 2F). Phosphorylation of IRF3 was observed in both SARS-CoV-2 infected and poly(I:C) treated cells relative to untreated cells, with similar levels of pIRF3-S386 observed following all infection and treatment conditions (Figure 2F).

### SARS-CoV-2 infection is unable to inhibit downstream type I IFN signaling

SARS-CoV and MERS-CoV proteins can also block downstream IFN signaling to restrict the production of ISGs (de Wit et al., 2016). To evaluate if SARS-CoV-2 can inhibit type I IFN signaling in response to exogenous IFNβ treatment, we infected Calu-3 cells for 12 hours at a MOI of 1 and stimulated these cells with recombinant human IFNβ for 6 hours. We monitored gene expression levels of *IRF7* and *IFIT1* in these cells by qPCR. Validation of the antiviral efficacy of our recombinant IFNβ1 was carried out in human fibroblast (THF) cells that were pretreated with IFNβ1, followed by RNA and DNA virus infections. Pre-treatment of THF cells with recombinant IFNβ1 inhibited the replication of herpes simplex virus (HSV), vesicular stomatitis virus (VSV) and H1N1 in a dose-dependent manner (see supplementary Figure S3E).

SARS-CoV-2 genome levels were significantly higher in infected cells relative to mock infected cells (Figure 3A). Although SARS-CoV-2 UpE levels displayed a lower trend in SARS-CoV-2 infected + IFNβ treated cells relative to SARS-CoV-2 infected only cells, UpE levels were not significantly different after 6 hours of IFNβ treatment (Figure 3A). Exogenous IFNβ treatment significantly upregulated transcript levels of *IRF7* and *IFIT1* relative to mock treated Calu-3 cells (Figures 3B and 3C). Consistent with our RNA-seq data, SARS-CoV-2 infection induced mild but significant levels of *IRF7* and *IFIT1* transcripts relative to mock infected cells (Figures 3B and 3C). IFNβ-mediated induction of *IRF7* and *IFIT1* was not dampened by SARS-CoV-2 infection (Figures 3B and 3C). Upregulation of *IRF7* and *IFIT1* transcripts by H1N1 was comparable to gene expression levels in IFNβ and H1N1+IFNβ treated cells (see supplementary Figure S3C).

**Figure 3.**
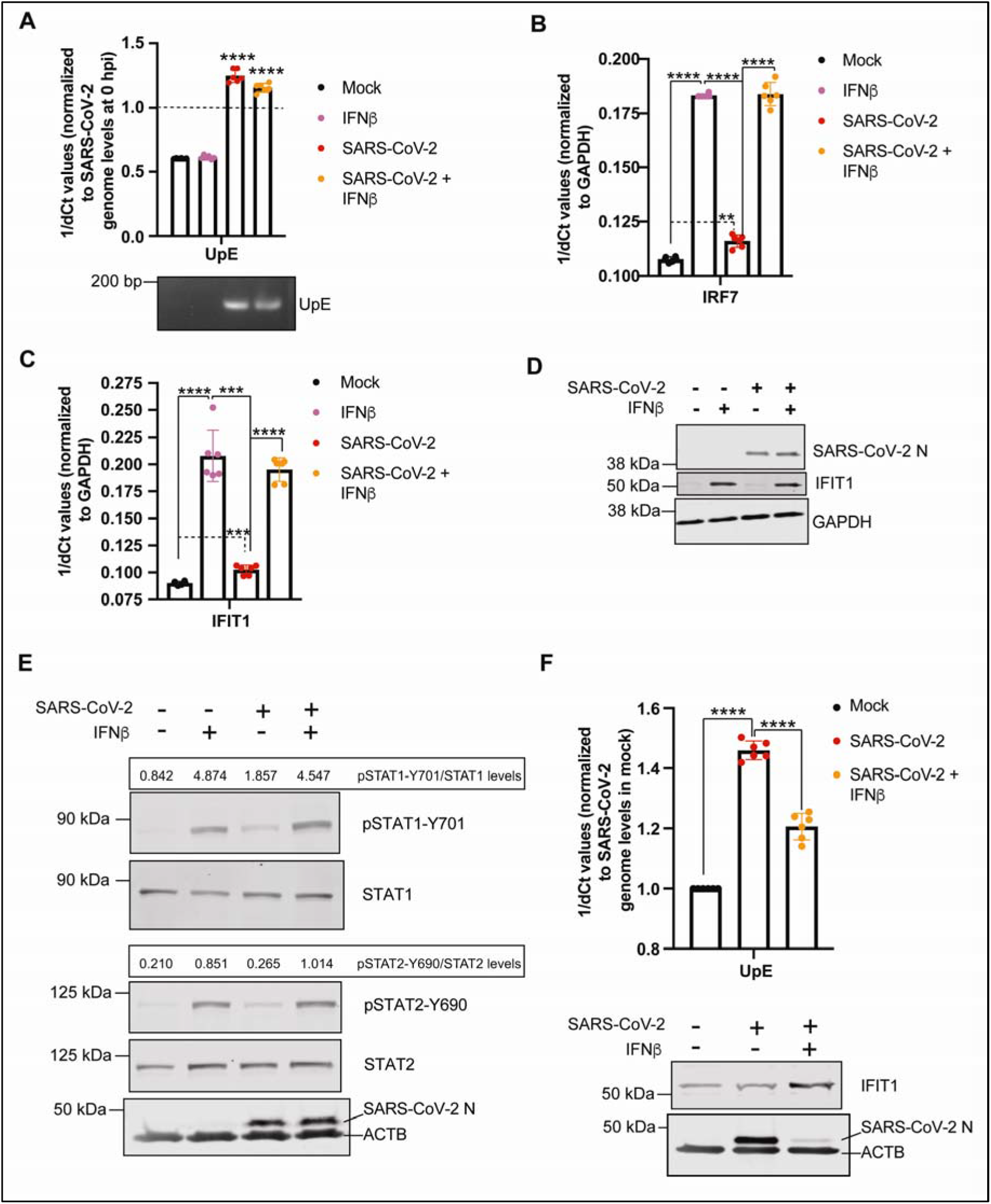
SARS-CoV-2 is unable to inhibit type I IFN signaling. To determine if SARS-CoV-2 can inhibit IFNβ-mediated stimulation of ISGs, such as IFIT1, Calu-3 cells were infected with SARS-CoV-2 at an MOI of 1 for 12 hours, following which cells were mock treated or treated with 2 mg/ml recombinant IFNβ. Mock infected and mock treated cells served as controls. Transcript levels were quantified using qPCR and protein expression was observed using immunoblots. (**A**) SARS-CoV-2 genome (UpE) levels in Calu-3 cells infected with SARS-CoV-2 or mock infected for 12 hours, and treated with recombinant IFNβ or mock treated for 6 hours (n = 6). 1/dCT values are represented after normalizing Ct values for SARS-CoV-2 genome levels at 18 hpi with Ct values observed at 0 hpi (immediately after removal of virus inoculum). Gel (below): UpE qPCR amplicons were also visualized on an agarose gel. (**B**) *IRF7* transcript levels in Calu-3 cells that were infected with SARS-CoV-2 or mock infected for 12 hours. Twelve hpi, cells were either treated with recombinant IFNβ or mock treated for 6 hours. *IRF7* transcript levels were normalized to *GAPDH* transcript levels (n = 6). (**C**) *IFIT1* transcript levels in Calu-3 cells that were infected with SARS-CoV-2 or mock infected for 12 hours. Twelve hpi, cells were either treated with recombinant IFNβ or mock treated for 6 hours. *IFIT1* transcript levels were normalized to *GAPDH* transcript levels (n = 6). (**D**) SARS-CoV-2 N, IFIT1 and GAPDH protein expression in Calu-3 cells that were infected with SARS-CoV-2 or mock infected for 12 hours. Twelve hpi, cells were either treated with recombinant IFNβ or mock treated for 6 hours (n = 3). (**E**) pSTAT1-Y701, STAT1, pSTAT2-Y690, STAT2, SARS-CoV-2 N and ACTB protein expression in Calu-3 cells that were infected with SARS-CoV-2 or mock infected for 24 hours. Twenty-four hpi, cells were either treated with recombinant IFNβ or mock treated for 30 minutes (n = 3). (**F**) SARS-CoV-2 genome (UpE) levels in Calu-3 cells infected with SARS-CoV-2 or mock infected for 1 hour followed by treatment with recombinant IFNβ or mock treatment for 72 hours (n = 6). 1/dCT values are represented after normalizing Ct values for SARS-CoV-2 genome levels in infected cells (with or without recombinant IFNβ treatment) with Ct values observed in mock infected cells. Blot (below): IFIT1, SARS-CoV-2 N and GAPDH protein expression in Calu-3 cells that were infected with SARS-CoV-2 or mock infected for 1 hr, followed by treatment with recombinant IFNβ or mock treatment for 72 hours (n = 3). Data are represented as mean ± SD, n = 3 or 6, *p*\*\*<0.01, ***<0.001 and ****<0.0001 (Student’s t test). Ct, cycle threshold. pSTAT1-Y701 and pSTAT2-Y690 protein expression levels are expressed as ratios of pSTAT1-Y701/STAT1 and pSTAT2-Y690/STAT2 levels, respectively. Blots were quantified using Image Studio (Li-COR) (n = 3). Ct, cycle threshold. See also supplementary Figures S3 and S4.

To validate our transcriptional responses, we repeated our experiments with exogenous IFNβ treatment and determined if SARS-CoV-2 can inhibit type I IFN-mediated upregulation of IFIT1 at the protein level. SARS-CoV-2 infection alone failed to induce detectable levels of IFIT1 at 12 hpi (Figure 3D). IFNβ treatment with or without prior 12 hours of SARS-CoV-2 infection induced robust expression of IFIT1 (Figure 3D). We confirmed SARS-CoV-2 infection in these cells by immunoblotting for N protein (Figure 3D). In comparison, H1N1 infection induced high levels of IFIT1 at 12 hpi, which was comparable to levels observed with IFNβ treatment (see supplementary Figure S3D). Additionally, similar to SARS-CoV-2 (Figure 3D), H1N1 infection failed to inhibit IFNβ-mediated upregulation of IFIT1 (see supplementary Figure S3D).

Binding of IFNs to their receptors activates a series of downstream signaling events, which involves phosphorylation of STAT1 at tyrosine 701 (pSTAT1-Y701) and STAT2 at tyrosine 690 (pSTAT2-Y690) (Pilz et al., 2003; Steen and Gamero, 2013). To determine if SARS-CoV-2 can inhibit phosphorylation of STAT1 and STAT2 proteins, we infected Calu-3 cells with SARS-CoV-2 for 24 hours followed by 30 minutes of stimulation with or without recombinant IFNβ. SARS-CoV-2 infection alone induced mild pSTAT1-Y701 and pSTAT2-Y690 levels relative to mock infected cells, albeit lower than levels observed in exogenous IFNβ treated cells (Figure 3E). Importantly, SARS-CoV-2 infection was unable to inhibit pSTAT1-Y701 and pSTAT2-Y690 levels in cells treated with IFNβ (Figure 3E).

To determine if exogenous IFNβ treatment can inhibit SARS-CoV-2 replication, we infected Calu-3 cells for 1 hour, following which we either mock treated or treated the cells with recombinant IFNβ for 72 hours. Exogenous IFNβ treatment reduced SARS-CoV-2 genome (UpE) and N protein levels in these cells (Figure 3F), consistent with an increase in IFIT1 levels (Figure 3F and see supplementary Figure S4).

### Serum cytokine levels vary in moderate and severe cases of COVID-19

To evaluate type I IFN and other infection-associated cytokines in COVID-19 patients, we analyzed acute sera (<21 days from symptom onset) from 20 COVID-19 positive patients, of whom 10 were categorized as ‘moderate’ cases requiring hospital admission, but not admission to intensive care unit (ICU). The remaining 10 samples were from ‘severe’ cases that required ICU admission. For severe cases, 6/10 patients died, and 10/10 moderate cases were discharged (see supplementary Table S4). We also included sera from 5 healthy, uninfected individuals. Sera from moderate cases of COVID-19 displayed significantly higher levels of platelet-derived growth factor AA (PDGF-AA) and PDGF-AB/BB relative to uninfected individuals (Figure 4). Patients with severe COVID-19 displayed significantly higher levels of PDGF-AA, PDGF-AB/BB, GROα (CXCL-1), CXCL-9, MIP-1β and vascular endothelial growth factor A (VEGF-A) relative to healthy individuals (Figure 4). Additionally, severe cases of COVID-19 displayed an increasing trend for levels of Interleukin-6 (IL-6), IL-5, macrophage colony-stimulating factor 1 (M-CSF), IL-8, tumor necrosis factor α (TNFα), TNFβ and granulocyte colony-stimulating factor 1 (G-CSF) relative to healthy individuals and moderate cases of COVID-19. Moderate cases of COVID-19 displayed an increasing trend for levels of IFN-α2 and IL-10 relative to healthy individuals and severe cases of COVID-19 (Figure 4 and see supplementary Tables S4 and S5). In addition, both moderate and severe cases of COVID-19 displayed an increasing trend for IL7 and IP-10 relative to healthy controls, although the data were not significant due to wide within patient variation in acute serum samples.

**Figure 4.**
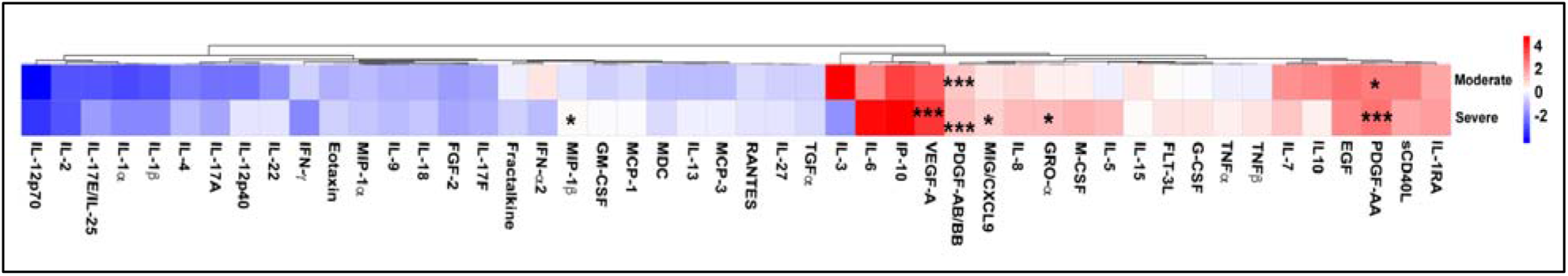
Cytokine protein levels in sera from moderate and severe cases of COVID-19 relative to healthy controls. To determine protein levels of cytokines in sera from moderate and severe cases of COVID-19 relative to healthy controls, we analyzed protein levels in sera using a 48-plex human cytokine and chemokine array. Mean log_2_ fold-change in serum cytokine protein levels in patients with moderate (n = 10) or severe (n = 10) case of COVID-19, relative to levels in healthy donors (n =5) are represented here. Data are represented as mean ± SD, n = 5 for healthy controls, and n = 10 each for moderate or severe cases of COVID-19, *p*\*<0.05, *p*\*\*<0.01 and ***<0.001 (Student’s t tests with Benjamini-Hochberg multiple testing correction). See also supplementary Tables S4 and S5.

## Discussion

SARS-CoV-2 emerged in December 2019 and has since caused a global pandemic of COVID-19 (Dong et al., 2020; Zhou et al., 2020a). Clinical observations and emerging data from *in vitro* and *in vivo* studies have demonstrated the ability of SARS-CoV-2 to induce type I IFNs (Blanco-Melo et al., 2020). However, a recent review summarized studies that suggest that antiviral IFN responses are dampened in COVID-19 patients (Acharya et al., 2020). Emerging data also suggest that timing and extent of interferon production is likely associated with manifestation of disease severity (Zhou et al., 2020b). In spite of some progress in understanding how SARS-CoV-2 activates antiviral responses, mechanistic studies into SARS-CoV-2-mediated modulation of human type I IFN responses are largely lacking. To understand SARS-CoV-2 infection-induced pathogenesis during the clinical course of COVID-19, it is imperative that we understand if and how SARS-CoV-2 interacts with type I IFN responses. These observations can be leveraged to develop drug candidates and inform ongoing drug trials, including trials that involve type I and III IFNs.

In this study, a robust time-series RNA-seq analysis of poly(A)-enriched RNA allowed us to map the progression of SARS-CoV-2 replication and transcription in Calu-3 cells using a clinical isolate (Banerjee et al., 2020a) of the virus. As observed with other coronaviruses (Fehr and Perlman, 2015; Lai, 1990; Perlman and Netland, 2009), SARS-CoV-2 replicated and transcribed sub-genomic RNA and mRNA in a directional manner (Figures 1A and 1B). Thus, our data show that SARS-CoV-2 replication strategy is consistent with other coronaviruses. Furthermore, our data demonstrate that Calu-3 cells support SARS-CoV-2 replication and that these cells represent a good *in vitro* model to study SARS-CoV-2-host interactions.

A recent study demonstrated the inability of SARS-CoV-2 to stimulate robust expression of type I IFNs in human lung cells (A549) that were genetically engineered to express angiotensin-converting enzyme 2 (ACE2), a cellular receptor for SARS-CoV-2 (Blanco-Melo et al., 2020); however, the mechanisms behind this dampened type I IFN response were not determined. Low SARS-CoV-2-induced type I IFN responses may be associated with (1) the virus’ ability to mask the detection of viral RNA by cellular PRRs and/or (2) the ability of viral proteins to inactivate cellular mechanisms involved in type I IFN induction (Shin et al., 2020). Data from our studies show that wildtype SARS-CoV-2 infection is capable of inducing a type I IFN response in human airway epithelial cells, including upregulation of IFN expression (Figures 1C, 1E and 1G) and downstream IFN signaling markers (Figures 1C, 1F, 1H, 1I and supplementary Figure S4). Our observations corroborate data from Lei *et al.’s* recent study where the authors demonstrated that SARS-CoV-2 infection of Calu-3 cells is capable of upregulating type I IFN responses (Lei et al., 2020).

Although SARS-CoV-2 infection is capable of inducing IFN responses, expression of *IFNβ*, *IFIT1* and *IRF7* is significantly lower relative to a potent synthetic inducer (eg. polyI:C) of IFNβ (Figures 2B-D). In comparison to H1N1 infection in Calu-3 cells, SARS-CoV-2 infection induces lower levels of *IFNβ*, *IFIT1* and *IRF7* transcripts (Figure 2B-D and supplementary Figure S3B). However, *IFNβ*, *IFIT1* and *IRF7* transcript levels remain significantly higher in SARS-CoV-2 infected cells relative to mock infected cells (Figures 2B-D). The physiological relevance of an existing, but dampened type I IFN response to SARS-CoV-2 remains to be identified. Emerging data suggest that prolonged and high levels of type I IFNs correlate with COVID-19 disease severity (Lucas et al., 2020). Thus, a dampened, yet protective early type I IFN response against SARS-CoV-2 may in fact be beneficial for humans (Park and Iwasaki, 2020). However, questions remain about how a low type I IFN response against SARS-CoV-2 could play a protective role during infection. One possibility would be that low levels of type I IFN production is sufficient to control SARS-CoV-2 replication, especially if the virus is unable to block downstream IFN signaling. This may explain the large number of asymptomatic cases of SARS-CoV-2 where an early IFN response may control virus replication and disease progression. Indeed, in one study, type I IFN (IFNα) levels were higher in asymptomatic cases relative to symptomatic cases (n=37) (Long et al., 2020). Further studies are required to identify regulatory mechanisms behind the protective role of a controlled and early IFN response during SARS-CoV-2 infection vs. the damaging long-term IFN response observed in severe cases of COVID-19.

SARS-CoV was shown in one study to inhibit poly(I:C)-mediated upregulation of IFNβ (Lu et al., 2011). However, our data show that infection with SARS-CoV-2 is unable to efficiently limit poly(I:C)-mediated upregulation of *IFNβ* transcripts and downstream ISGs, such as *IFIT1* and *IRF7* (Figures 2B-D). The limitation of Lu *et al.’s* study with SARS-CoV and other recent studies with SARS-CoV-2 is the use of protein over-expression models to identify host response modulating capabilities of viral proteins (Gordon et al., 2020; Jiang et al., 2020; Lei et al., 2020; Xia et al., 2020). The lack of wildtype virus infection does not represent a realistic scenario, where the dynamics of virus replication (PAMP generation) and protein translation (modulators of type I IFN response) would affect the overall antiviral outcome. Indeed, in Lei *et al’s* recent study, wildtype SARS-CoV-2 infection significantly upregulated transcript levels of *IFNβ* and *IFIT1*; however, ectopic expression of individual proteins led to the identification of 8 SARS-CoV-2 proteins that could inhibit the activation of the IFNβ promoter (Lei et al., 2020). The physiological relevance of over-expressing SARS-CoV-2 proteins that can block type I IFN responses remains to be validated, but it is unlikely that SARS-CoV-2 proteins will be selectively upregulated to such high amounts during the natural course of infection, while limiting the generation of stimulatory RNA molecules during virus replication. Similarly, ectopic expression of the NS1 protein from H1N1 influenza A virus strongly inhibits type I IFN responses (Jia et al., 2010; Kochs et al., 2007); however, in contrast, infection with wildtype H1N1 virus induces a type I IFN response (Jewell et al., 2007) (see supplementary Figures S3B and S3C). Thus, the physiological relevance of observations made by over-expressing viral proteins is debatable and for translatability, it is critical to understand IFN production and modulation using wildtype virus infection. In our study, poly(I:C) transfection + SARS-CoV-2 infection induced higher levels of ISG (*IFIT1*) transcripts relative to poly(I:C) alone, indicating that wildtype infection partially augments poly(I:C)-mediated upregulation of type I IFN signaling (Figures 2C and 2D) in Calu-3 cells. Thus, it is important to identify the kinetics and landscape of virus infection, transcription and translation, and how that may regulate human type I IFN responses.

Coronaviruses, including highly pathogenic SARS-CoV, MERS-CoV and porcine epidemic diarrhea virus (PEDV) have evolved proteins that can efficiently inhibit type I IFN responses (Chen et al., 2014; Ding et al., 2014; Lu et al., 2011; Lui et al., 2016; Niemeyer et al., 2013; Siu et al., 2014; Xing et al., 2013; Yang et al., 2013). Our data show that SARS-CoV-2 infection does not modulate poly(I:C)-mediated phosphorylation of TBK1 and IRF3 (Figure 2F) and downstream upregulation of transcript and protein levels of *IFIT1* (Figures 2C and 2E). In spite of observing statistically significant upregulation of type I IFNs and ISGs at 12 hpi with SARS-CoV-2 (Figure 1), in preliminary studies we were unable to observe detectable levels of pIRF3-S386 prior to accumulation of antiviral mRNAs. We have previously shown that antiviral responses can be induced in the absence of prototypic markers of IRF3 activation such as dimerization and hyperphosphorylation, even when IRF3 was shown to be essential (Noyce et al., 2009). The simplest interpretation is that early activation of IRF3-mediated IFN responses requires low (or even undetectable) levels of pIRF3-S386, which accumulate to detectable levels over time (Figure 2F).

SARS-CoV and MERS-CoV can inhibit phosphorylation and activation of STAT1 and STAT2, which blocks global IFN-induced antiviral responses (de Wit et al., 2016). Our data demonstrate that SARS-CoV-2 infection induces phosphorylation of STAT1 and STAT2 (Figure 3E), along with upregulation of ISGs, such as *IRF7* and *IFIT1* (Figures 3B and 3C). In addition, SARS-CoV-2 infection is unable to inhibit the activation of STAT1 and STAT2 by exogenous type I IFN (Figure 3E), along with the expression of downstream ISGs, such as *IRF7* and *IFIT1* (Figures 3B and 3C and see supplementary Figure S4). Although SARS-CoV-2 infection alone induced low levels of type I IFN (Figures 1E and 2B), it was sufficient to activate STAT proteins (Figure 3E) and downstream ISG expression (Figures 2C, 2D, 3B and 3C; see supplementary Figures S1 and S2). Thus, the dampened ability of SARS-CoV-2 to block downstream type I IFN responses compared to other zoonotic CoVs extends support to our hypothesis that the pathogenic consequences of a dampened type I IFN response may be largely negated by the sensitivity of SARS-CoV-2 to this response. Indeed, in our studies, exogenous type I IFN (IFNβ1) treatment significantly reduced SARS-CoV-2 replication in human airway epithelial cells (Figure 3F). Our data provide promising support for ongoing clinical trials that include type I IFN treatment.

Recent studies have shown that COVID-19 patients mount a dysregulated immune response, which is associated with a poor clinical outcome (Lucas et al., 2020). In our study, we observed that patients with moderate or severe case of COVID-19 had elevated serum levels of growth factors PDGF-AA and PDGF-AB/BB relative to healthy controls (Figure 4 and supplementary Tables 4 and 5). The role of PDGFs in driving disease pathology has been described previously (Andrae et al., 2008) and therapeutic use of PDGF antagonists have also been recommended (Grimminger and Schermuly, 2010; Sadiq et al., 2015). PDGF-BB has also been introduced in the clinic as a wound-healing therapy (Yamakawa and Hayashida, 2019). The physiological impact of elevated PDGF levels and cellular factors that regulate the expression of PDGF in COVID-19 patients remains to be understood.

Sera from patients with moderate case of COVID-19 contained higher levels of IL-10 relative to severe cases, which is suggestive of an anti-inflammatory response (Couper et al., 2008; Pripp and Stanisic, 2014) (Figure 4 and supplementary Tables 4 and 5). On the contrary, sera from patients with severe case of COVID-19 displayed a higher trend for levels of IL-6, IL-8 and TNFα relative to moderate cases, which is suggestive of a pro-inflammatory response (Figure 4 and supplementary Tables 4 and 5) (Lucas et al., 2020; Mandel et al., 2020; Pripp and Stanisic, 2014). Observations from our study (Figure 4), along with other recent reports (Long et al., 2020; Lucas et al., 2020) warrant further investigations into mechanistic regulation of pro- and anti-inflammatory processes in SARS-CoV-2 infected human airway cells. Identifying regulatory proteins, such as transcription factors that contribute to a pro-inflammatory cytokine response in SARS-CoV-2 infected cells will inform the selection and utilization of anti-inflammatory drugs.

Moderate cases of COVID-19 demonstrated an increasing trend for type I IFN (IFN-α2) relative to severe cases and healthy controls (Figure 4). In a separate study, IFN-α levels were also higher in asymptomatic patients relative to symptomatic COVID-19 patients (Long et al., 2020). The presence of type I IFN in moderate cases of COVID-19 in our study, along with a recent study by Lucas *et al.* suggest that SARS-CoV-2 infection is capable of inducing a type I IFN response *in vivo*; however, emerging clinical data suggest that the extent and duration of type I IFN response may dictate the clinical course of COVID-19 (Hadjadj et al., 2020b; Lucas et al., 2020). In our study, sera from COVID-19 patients were collected at admission (all <21 days post symptom onset). Early induction of IFN-α2 in moderate cases of COVID-19 may provide an antiviral advantage. We were unable to detect IFN-α2 in severe cases at this sampling time. Additional studies with later samples from severe COVID-19 patients will identify if there is a late and prolonged induction of type I IFNs as reported recently by Lucas *et al.* (Lucas et al., 2020). In spite of recent progress in understanding type I IFN responses in COVID-19 patients, factors associated with early or delayed and short-acting vs. prolonged type I IFN induction in COVID-19 patients remains to be understood.

In conclusion, our study demonstrates that SARS-CoV-2 is a weak stimulator of type I IFN production in infected human airway epithelial cells, relative to poly(I:C) or H1N1 virus infection. However, our data suggest that low levels of type I IFN response in SARS-CoV-2 infected cells is sufficient to activate downstream expression of antiviral ISGs. In addition, our data demonstrate that SARS-CoV-2 is unable to inhibit downstream IFN responses that are mediated by STAT proteins, which is promising for the development of type I IFNs as treatment or post-exposure prophylactics. Clinical trials for combination IFNβ therapy against MERS-CoV are currently ongoing (Arabi et al., 2020). IFNβ, in combination with lopinavir-ritonavir and ribavirin has been used with promising results in COVID-19 patients (Hung et al., 2020).

Nebulized IFNβ is part of the standard of care for COVID-19 patients in China (Xu et al., 2020). Importantly, our data with wildtype virus infection appears to be in contrast to functions observed for inhibition of IFN induction and signaling via protein over-expression studies. Thus, our study highlights the dynamic nature of virus-host interaction during the course of SARS-CoV-2 infection and demonstrates that SARS-CoV-2 has not evolved to efficiently block human antiviral interferon responses.

## Acknowledgements

This study was supported by a Canadian Institutes of Health Research (CIHR) COVID-19 rapid response grant to principal applicant K.M. and Co-Applicants A.B., A.G.M., M.S.M. and S.M. A.B. was funded by the Natural Sciences and Engineering Research Council of Canada (NSERC). Computer resources were in part supplied by the McMaster Service Lab and Repository computing cluster, funded in part by grants to A.G.M. from the Canadian Foundation for Innovation. J.A.H. is supported by the Canada Research Chairs Program and an Ontario Early Career Researcher Award. M.S.M. is supported by a CIHR COVID-19 rapid response grant, a CIHR New Investigator Award and an Ontario Early Researcher Award. ATI is supported by a Zhejiang University scientific research fund for COVID-19 prevention and control.

The graphical abstract was created using BioRender.com.

## Author Contributions

Conceptualization, A.B., D.R., P.B., H.M., A.G.M., A.C.D., J.A.A. and K.M.; Methodology, A.B., N.E-.S., P.B., D.R., H.M., K.B., M.R.D., S.A., M.K. and L.Y.; Formal analysis A.B., N.E-S., P.B., D.R., H.M., J.A.A, K.B., B.J.-M.T., and M.K.; Reagents, J.A.H., A.J.M, and M.S.M.; Funding acquisition, A.B., M.O., J.A.H., R.K., T.D.C., M.S.M., B.W., S.M., A.G.M, A.C.D., and K.M.; Writing – reviewing and editing A.B., N.E-.S., P.B., D.R., H.M., J.A.A., K.B., J.A.H, A.T.I, A.G.M., A.C.D. and K.M. All authors reviewed the final manuscript. Supervision, T.D.C, M.S.M., B.W., S.M., A.G.M, A.C.D and K.M.

## Declaration of Interests

The authors declare no competing interests.

### Transparent Methods

#### Contact for Reagent and Resource Sharing

Dr. Karen Mossman and Dr. Arinjay Banerjee.

#### Computational Analysis

##### Transcript quantification and differential expression analysis

Sequence read quality was checked with FastQC (https://www.bioinformatics.babraham.ac.uk/projects/fastqc/), with reads subsequently aligned to the human reference transcriptome (GRCh37.67) obtained from the ENSEMBL database (Hunt et al., 2018), indexed using the ‘index’ function of Salmon (version 0.14.0) (Patro et al., 2017) with a k-mer size of 31. Alignment was performed using the Salmon ‘quant’ function with the following parameters: “-l A --numBootstraps 100 --gcBias --validateMappings”. All other parameters were left to defaults. Salmon quantification files were imported into R (version 3.6.1) (RCoreTeam, 2017) using the tximport library (version 1.14.0) (Soneson et al., 2015) with the ‘type’ option set to ‘salmon’. Transcript counts were summarized at the gene-level using the corresponding transcriptome GTF file mappings obtained from ENSEMBL. Count data was subsequently loaded into DESeq2 (version 1.26.0) (Love et al., 2014) using the ‘DESeqDataSetFromTximport’ function. In order to determine time/treatment dependent expression of genes, count data was normalized using the ‘estimateSizeFactors’ function using the default ‘median ratio method’ and output using the ‘counts’ function with the ‘normalized’ option.

For subsequent differential-expression analysis, a low-count filter was applied prior to normalization, wherein a gene must have had a count greater than 5 in at least three samples in order to be retained. Using all samples, this resulted in the removal of 12,980 genes for a final set of 15,760 used. Principal Component Analysis (PCA) of samples across genes was performed using the ‘vst’ function in DESeq2 (default settings) and was subsequently plotted with the ggplot2 package in R (Wickham, 2009). Differential expression analyses were carried out with three designs: (a) the difference between infection/control status across all timepoints, (b) considering the effects of post-infection time (i.e. the interaction term between time and infection status) and (c) the difference between infection/control status at individual timepoints. (a) and (b) were performed using the ‘DESeq’ function of DESeq2 using all samples, with results subsequently summarized using the ‘results’ function with the ‘alpha’ parameter set to 0.05; *p*-values were adjusted using the Benjamini-Hochberg FDR method (Benjamini and Hochberg, 1995), with differentially expressed genes filtered for those falling below an adjusted *p*-value of 0.05. For (c), infected/mock samples were subset to individual timepoints, with differential expression calculated using DESeq as described above. Additionally, given the smaller number of samples at individual time-points, differential-expression analysis was also performed with relaxation of the low-count filter described above, with results and p-value adjustments performed as above.

##### Viral transcript quantification

Paired-end sequencing reads were mapped to CDS regions of the SARS-CoV-2 genomic sequence (Assembly ASM985889v3 - GCF_009858895.2) obtained from NCBI, indexed using the ‘index’ function of Salmon (version 0.14.0) (Patro et al., 2017) with a k-mer size of 31. Subsequently, reads were aligned using the Salmon ‘quant’ function with the following parameters: “-l A --numBootstraps 100 --gcBias --validateMappings”. All other parameters were left to defaults. Salmon quantification files were imported into R (version 3.6.1) (RCoreTeam, 2017) using the tximport library (version 1.14.0) (Soneson et al., 2015) with the ‘type’ option set to ‘salmon’. All other parameters were set to default. Transcripts were mapped to their corresponding gene products via GTF files obtained from NCBI. Count data was subsequently loaded into DESeq2 (version 1.26.0) (Love et al., 2014) using the ‘DESeqDataSetFromTximport’ function. Principal Component Analysis (PCA) of samples across viral genes was performed using the ‘vst’ function in DESeq2 (default settings) and was subsequently plotted with the ggplot2 package in R (42) (Figure 1A). As viral transcript levels increased over time post-infection, we first converted non-normalized transcript counts to a log_2_ scale, and subsequently compared these across time-points (Figure 1B and supplementary Table S1). To look at the changes in the expression of viral transcripts relative to total viral expression as a function of post-infection time, normalized transcript counts were used to perform differential-expression analysis with DESeq2. Results and *p*-value adjustments were performed as described above.

In order to compare host/viral expression patterns, normalized transcript counts from infected samples were compared with either normalized or non-normalized viral transcript counts (from the same sample) across time-points. For each viral transcript (n = 12), all host genes (n = 15,760, after filtering described above) were tested for correlated expression changes across matched infected samples (n = 18, across 5 time-points) using Pearson’s correlation coefficient (via the cor.test function in R). Correlation test *p*-values were adjusted across all-by-all comparisons using the Benjamini-Hochberg FDR method, and gene-transcript pairs at adjusted *p*< 0.05 were retained. To account for possible effects of cellular response to plate incubation, viral transcript abundance was averaged at each time-point and compared to host transcript abundance similarly averaged at each time-point for non-infected samples; correlation testing was done all-by-all for n = 5 data-points. Host genes that correlated with viral transcription in mock samples across time were removed from subsequent analyses; to increase stringency, mock correlation was defined using un-adjusted *p*< 0.05. Host genes were sorted by correlation coefficient (with any given viral transcript), with the top 100 unique genes retained for visualization. Normalized host transcript counts were z-score transformed per-gene using the ‘scale’ function in R, with normalized/un-normalized viral transcript counts similarly transformed per-transcript. Resulting z-score expression heatmaps were generated using the ComplexHeatmap library in R (version 2.2.0) (Gu et al., 2016). Heatmaps were generated for normalized/un-normalized viral transcript counts, given the different information revealed by absolute and relative viral expression patterns.

##### Viral genome mapping

Paired-end RNA-seq reads were filtered for quality control with Trim Galore! (version 0.6.4_dev) (Krueger, 2019) and mapped to the SARS-CoV-2 reference sequence (NC_045512.2) with the Burrow-Wheeler Aligner (Li and Durbin, 2009), using the BWA-MEM algorithm (Li, 2013). Output SAM files were sorted and compressed into BAM files using Samtools (version 1.10) (Li et al., 2009). Read coverage visualization was performed from within the R statistical environment (version 4.0.0) (RCoreTeam, 2017) using the “scanBam” function from the Rsamtools R package (version 1.32.0) to extract read coverage data and the ggplot2 R package (version 3.3.0) (Wickham, 2009) to plot read coverage histograms (using 300 bins across the SARS-CoV-2 sequence).

##### Cellular pathway enrichment analysis

To determine cellular pathways that were associated with differentially expressed genes (DEGs), the ActivePathways R (version 1.0.1) (Paczkowska et al., 2020) package was utilized to perform gene-set based pathway enrichment analysis. DEGs at each time point were treated as an independent set for enrichment analysis. Fisher’s combined probability test was used to enrich pathways after *p*-value adjustment using Holm-Bonferroni correction. Pathways of gene-set size less than 5 and greater than 1000 were excluded. Only pathways enriched at individual time-points were considered for downstream analysis; pathways enriched across combined timepoints as determined by ActivePathways Brown’s *p*-value merging method were filtered out.

Visualization of enriched pathways across timepoints was done using Cytoscape (version 3.8.0) (Shannon et al., 2003) and the EnrichmentMap plugin (version 3.2.1) (Merico et al., 2010), as outlined by Reimand *et al*. (Reimand et al., 2019). Up-to-date Gene-Matrix-Transposed (GMT) files containing information on pathways for the Gene Ontology (GO) Molecular Function (MF), GO Biological Process (BP) (The Gene Ontology, 2019) and REACTOME (Jassal et al., 2020) pathway databases were utilized with ActivePathways. Only pathways that were enriched at specific time points were considered. Bar plots displaying top ActivePathway GO terms and REACTOME enrichments for infection versus mock were plotted using the ggplot2 R package (version 3.2.1) for 1-, 2-, 3-, and 12-hour time points. Zero and 6-hour time points were omitted due to a lack of sufficient numbers of differentially expressed genes required for functional enrichment analysis.

#### Experimental Model

##### Cells and viruses

Vero E6 cells (African green monkey cells; ATCC) were maintained in Dulbecco’s modified Eagle’s media (DMEM) supplemented with 10% fetal bovine serum (FBS; Sigma-Aldrich), 1x L-Glutamine and Penicillin/Streptomycin (Pen/Strep; VWR). Calu-3 cells (human lung adenocarcinoma derived; ATCC) were cultured as previously mentioned (Aguiar et al., 2019). THF cells (human telomerase life-extended cells; from Dr. Victor DeFilippis’ lab) were cultured as previously mentioned (Banerjee et al., 2020b). *Drosophila* S2 cells (ThermoFisher Scientific) were cultured in Schneider’s *Drosophila* medium supplemented with 10% FBS (Sigma-Aldrich) as recommended by the manufacturer and cells were incubated at 28°C. Stocks of genetically engineered vesicular stomatitis virus (VSV-GFP) carrying a green fluorescent protein (GFP) cassette (Noyce et al., 2011) were stored at −80°C. H1N1 (A/Puerto Rico/8/1934 mNeon – 2A-HA) stocks were obtained from Dr. Matthew Miller’s laboratory.

HSV-GFP stocks were generated and maintained as mentioned previously (Minaker et al., 2005). Clinical isolate of SARS-CoV-2 (SARS-CoV-2/SB3) was propagated on Vero E6 cells and validated by next generation sequencing (Banerjee et al., 2020a). Virus stocks were thawed once and used for an experiment. A fresh vial was used for each experiment to avoid repeated freeze-thaws. VSV-GFP, HSV-GFP and H1N1 infections were performed at a multiplicity of infection (MOI) of 1. SARS-CoV-2 infections were performed at an MOI of 1 or 2. Experiments with SARS-CoV-2 were performed in a BSL3 laboratory and all procedures were approved by institutional biosafety committees at McMaster University and the University of Toronto.

#### Method Details

##### RNA-Seq

RNA was isolated from cells using RNeasy Mini kit (Qiagen). Sequencing was conducted at the McMaster Genomics Facility, Farncombe Institute at McMaster University. Sample quality was first assessed using a Bioanalyzer (Agilent), then enriched (NEBNext Poly(A) mRNA Magnetic Isolation Module; NEB). Library preparations were conducted (NEBNext Ultra II Directional RNA Library Prep Kit; NEB) and library fragment size distribution was verified (Agilent TapeSection D1000; Agilent). Libraries were quantified by qPCR, pooled in equimolar amounts, and qPCR and fragment size distribution verification were conducted again. Libraries were then sequenced on an Illumina HiSeq 1500 across 3 HiSeq Rapid v2 flow cells in 6 lanes (Illumina) using a paired-end, 2×50 bp configuration, with onboard cluster generation averaging 30.8M clusters per replicate (minimum 21.9M, maximum 46.0M).

##### Cytokine levels in COVID-19 patient sera

Acute patient sera (<21 days from symptom onset) were acquired from moderate (hospital admission, but no ICU admission) and severe (ICU admission) cases of COVID-19 in Toronto, Canada, along with samples from uninfected, healthy individuals (see supplementary Table S4). Sera were analyzed using a 48-plex human cytokine and chemokine array by the manufacturer (Evetechnologies). Samples with an observed cytokine concentration (pg/ml) below the limit of detection (OOR<) were floored to the lowest observed concentration for that cytokine. Average log_2_FC for moderate patients (n=10) vs healthy patients (n=5), and severe patients (n=10) vs healthy patients (n=5) were plotted using the pheatmap() R package (version 3.2.1) for all of the 48 cytokines. Cytokine expression levels were tested for significant differences via unpaired Student’s t tests with Benjamini-Hochberg multiple testing correction using the *stats* R package (version 3.6.1). Work with patient sera was approved by the Sunnybrook Research Institute Research Ethics Board (amendment to 149-1994, March 2, 2020) (Nasir et al., 2020).

##### Poly(I:C) transfection and IFNβ treatment

Calu-3 cells were mock transfected with 4 μl of lipofectamine 3000 (ThermoFisher Scientific) in Opti-MEM (ThermoFisher Scientific) only or transfected with 100 to 1000 ng of poly(I:C) (InvivoGen). Recombinant human IFNβ1 was generated using *Drosophila* Schneider 2 (S2) cells following manufacturer’s recommendation and by using ThermoFisher Scientific’s *Drosophila* Expression system (ThermoFisher Scientific). As a control, recombinant GFP was also generated using the same protocol and used for mock treated cells. For VSV-GFP, HSV-GFP and H1N1-mNeon infections, cells were treated with increasing concentrations of IFNβ1 or GFP (control). SARS-CoV-2 infected cells were treated with 2 mg/ml of IFNβ1 or GFP.

##### Quantitative PCR

Calu-3 cells were seeded at a density of 3 × 10^5^ cells/well in 12-well plates. Cells were infected with SARS-CoV-2 for 12 hours. Twelve hours post incubation, mock infected or infected cells were mock stimulated or stimulated with poly(I:C) or IFNβ for 6 hours. RNA extraction was performed using RNeasy^®^ Mini Kit (Qiagen) according to manufacturer’s protocol 6 hours post poly(I:C) tranfection or . 200 ng of purified RNA was reverse transcribed using iScript™ gDNA Clear cDNA Synthesis Kit (Bio-Rad). Quantitative PCR reactions were performed with TaqMan™ Universal PCR Master Mix (ThermoFisher Scientific) using pre-designed Taqman gene expression assays (ThermoFisher Scientific) for *IFNβ1* (catalog no. #4331182), *IRF7* (catalog no. #4331182), *IFIT1* (catalog no. #4331182) and *GAPDH* (catalog no. #4331182) according to manufacturer’s protocol. Relative mRNA expression was normalized to *GAPDH* and presented as 1/ΔCt. To quantify SARS-CoV-2 genome levels, primers were designed to amplify a region (UpE) between *ORF3a* and *E* genes. Primer sequences used were SARS2 UpE F – ATTGTTGATGAGCCTGAAG and SARS2 UpE R–TTCGTACTCATCAGCTTG. qPCR to determine UpE levels was performed using SsoFast EvaGreen supermix (Bio-Rad) as previously described (Banerjee et al., 2017).

##### Immunoblots

Calu-3 cells were seeded at a density of 3 × 10^5^ cells/well in 12-well plates. Cells were infected with SARS-CoV-2 at an MOI of 1. Control cells were sham infected. Twelve to twenty-four hours post incubation, cells were transfected or treated with poly(I:C) or IFNβ, respectively for indicated times. Cell lysates were harvested for immunoblots and analyzed on reducing gels as mentioned previously (Banerjee et al., 2020b). Briefly, samples were denatured in a reducing sample buffer and analyzed on a reducing gel. Proteins were blotted from the gel onto polyvinylidene difluoride (PVDF) membranes (Immobilon, EMD Millipore) and detected using primary and secondary antibodies. Primary antibodies used were: 1:1000 mouse anti-GAPDH (EMD Millipore; Catalogue number: AB2302; RRID: AB_10615768), 1:1000 mouse anti-SARS/SARS-CoV-2 N (ThermoFisher Scientific; Catalogue number: MA5-29981; RRID: AB_2785780, 1:1000 rabbit anti-IFIT1 (ThermoFisher Scientific; Catalogue number: PA3-848; RRID: AB_1958733), 1:1000 rabbit anti-beta-actin (Abcam; Catalogue number: ab8227; RRID:), 1:1000 rabbit anti-IRF3 (Abcam; Catalogue number: ab68481; RRID: AB_11155653), 1:1000 rabbit anti-pIRF3-S386 (Cell Signaling; Catalogue number: 4947; RRID: AB_823547), 1:1000 rabbit anti-TBK1 (Abcam; Catalogue number: ab40676; RRID: AB_776632), 1:1000 rabbit anti-pTBK1-S172 (Abcam; Catalogue number: ab109272; RRID: AB_10862438), 1:1000 rabbit anti-STAT1 (Cell Signaling; Catalogue number: 9172; RRID: AB_2198300), 1:1000 rabbit anti-pSTAT1-Y701 (Cell Signaling; Catalogue number: 9167; RRID: AB_561284), 1:1000 rabbit anti-STAT2 (Cell Signaling; Catalogue number: 72604; RRID: AB_2799824), 1:1000 rabbit anti-pSTAT2-Y690 (Cell Signaling; Catalogue number: 88410S; RRID: AB_2800123). Secondary antibodies used were: 1:5000 donkey anti-rabbit 800 (LI-COR Biosciences; Catalogue number: 926-32213; RRID: 621848) and 1:5000 goat anti-mouse 680 (LI-COR Biosciences; Catalogue number: 925-68070; RRID: AB_2651128). Blots were observed and imaged using Image Studio (LI-COR Biosciences) on the Odyssey CLx imaging system (LI-COR Biosciences).

##### Antiviral bioassay

THF cells were pre-treated or mock treated with recombinant human IFNβ, followed by VSV-GFP, HSV-GFP or H1N1-mNeon infection at an MOI of 1. Infected cells were incubated at 37°C for 1 hour with gentle rocking every 15 minutes. After 1 hour, virus inoculum was aspirated and Minimum Essential Medium (MEM) with Earle’s salts (Sigma) containing 2% FBS and 1% carboxymethyl cellulose (CMC; Sigma) was added on the cells. Cells were incubated for 19 hours at 37°C and green fluorescent protein (GFP) or mNeon levels were measured using a typhoon scanner (Amersham, Sigma).

#### Statistical Analysis

Statistical analyses for RNA-seq data were performed in R and are mentioned under the respective RNA-seq analyses sections. All other statistical calculations were performed in GraphPad Prism (version 8.4.2; www.graphpad.com) using two-tailed paired t-test. Significance values are indicated in the figures and figure legends. *p*\*<0.05, **<0.01, ***<0.001 and ****<0.0001.

#### Data Availability

The DESeq2 normalized transcript counts for all genes with RNA-Seq data, significant or otherwise, plus the raw sequencing FASTQ reads have been deposited into the Gene Expression Omnibus (GEO) database with NCBI GEO accession number GSE151513.

## Supplemental Items

### Tables

**Table S1.** Mean raw read counts for SARS-CoV-2 transcripts. INF, SARS-CoV-2 infected; H, hours post incubation; SD, standard deviation. Related to Figure 1.

| Mean<br>INF<br>0H | SD<br>INF<br>0H | Mean<br>INF<br>1H | SD<br>INF<br>1H | Mean<br>INF<br>2H | SD<br>INF<br>2H | Mean<br>INF<br>3H | SD<br>INF<br>3H | Mean<br>INF<br>6H | SD<br>INF<br>6H | Mean<br>INF<br>12H | SD<br>INF<br>12H | SARS-CoV-2<br>gene | Transcript |
| --- | --- | --- | --- | --- | --- | --- | --- | --- | --- | --- | --- | --- | --- |
| 257.67 | 38.59 | 285.33 | 56.13 | 243.67 | 39.25 | 278.00 | 23.00 | 12173.3 | 3006.93 | 25827.33 | 2054.93 | ORF1ab | lcl\|NC_045512.2<br>_cds_YP_00972<br>4389.1_1 |
| 1.00 | 1.73 | 0.00 | 0.00 | 0.33 | 0.58 | 0.00 | 0.00 | 33.67 | 6.81 | 1061.00 | 468.03 | ORF1a | lcl\|NC_045512.2<br>_cds_YP_00972<br>5295.1_2 |
| 500.67 | 94.52 | 491.33 | 86.19 | 378.00 | 61.39 | 521.67 | 49.69 | 19232.3 | 3952.46 | 26903.33 | 3860.82 | spike | lcl\|NC_045512.2<br>_cds_YP_00972<br>4390.1_3 |
| 173.67 | 24.99 | 172.33 | 43.68 | 127.33 | 17.16 | 203.33 | 26.50 | 9995.00 | 1736.00 | 13976.33 | 2233.55 | ORF3a | lcl\|NC_045512.2<br>_cds_YP_00972<br>4391.1_4 |
| 43.67 | 5.51 | 44.67 | 13.65 | 39.00 | 2.65 | 63.00 | 11.53 | 2903.33 | 485.15 | 4086.33 | 627.70 | envelope | lcl\|NC_045512.2<br>_cds_YP_00972<br>4392.1_5 |
| 199.67 | 27.02 | 196.00 | 37.32 | 162.33 | 28.87 | 298.67 | 19.60 | 22344.3 | 3354.18 | 31200.33 | 4915.23 | membrane | lcl\|NC_045512.2<br>_cds_YP_00972<br>4393.1_6 |
| 34.67 | 2.08 | 32.33 | 10.50 | 25.00 | 7.81 | 45.33 | 1.53 | 3508.00 | 509.12 | 4704.67 | 886.56 | ORF6 | lcl\|NC_045512.2<br>_cds_YP_00972<br>4394.1_7 |
| 107.33 | 19.50 | 102.33 | 23.35 | 94.00 | 22.61 | 173.67 | 34.00 | 14834.0 | 2357.53 | 21920.67 | 3441.71 | ORF7a | lcl\|NC_045512.2<br>_cds_YP_00972<br>4395.1_8 |
| 10.33 | 2.52 | 11.67 | 2.31 | 15.33 | 1.53 | 20.67 | 1.15 | 1516.33 | 241.00 | 2191.33 | 526.17 | ORF7b | lcl\|NC_045512.2<br>_cds_YP_00972<br>5318.1_9 |
| 109.33 | 22.19 | 107.00 | 27.22 | 98.00 | 21.70 | 189.00 | 14.00 | 14651.3 | 2136.80 | 21518.67 | 3992.04 | ORF8 | lcl\|NC_045512.2<br>_cds_YP_00972<br>4396.1_10 |
| 1251.00 | 230.97 | 1157.33 | 247.52 | 1067.67 | 144.58 | 2945.67 | 402.61 | 258553 | 34843.96 | 393221.67 | 62159.07 | nucleocapsid | lcl\|NC_045512.2<br>_cds_YP_00972<br>4397.2_11 |
| 112.33 | 27.57 | 97.00 | 22.52 | 94.67 | 10.69 | 250.00 | 19.00 | 18385.3 | 2239.71 | 27679.00 | 5406.01 | ORF10 | lcl\|NC_045512.2<br>_cds_YP_00972<br>5255.1_12 |

**Table S2.** Mean normalized read counts for differentially expressed IFN and ISG transcripts. H, hours post incubation; INF, SARS-CoV-2 infected; MOCK, mock infected; IFN, interferon; ISG, interferon stimulated genes. Related to Figure 1.

|  |  | 0H<br>INF<br>(N=3) | 0H<br>MOCK<br>(N=3) | 1H<br>INF<br>(N=3) | 1H<br>MOCK<br>(N=3) | 2H<br>INF<br>(N=3) | 2H<br>MOCK<br>(N=3) | 3H<br>INF<br>(N=3) | 3H<br>MOCK<br>(N=3) | 6H<br>INF<br>(N=3) | 6H<br>MOCK<br>(N=3) | 12H<br>INF<br>(N=3) | 12H<br>MOCK<br>(N=3) |
| --- | --- | --- | --- | --- | --- | --- | --- | --- | --- | --- | --- | --- | --- |
| IFNs | IFNB1 | 1.35 | 0.00 | 1.21 | 0.41 | 1.48 | 0.97 | 0.57 | 1.93 | 6.40 | 0.30 | 21.23 | 0.89 |
|  | IFNL1 | 3.49 | 3.45 | 2.20 | 4.80 | 4.93 | 5.46 | 3.17 | 1.90 | 7.00 | 2.66 | 15.07 | 0.73 |
|  | IFNL2 | 0.00 | 0.00 | 0.36 | 0.96 | 4.11 | 0.35 | 0.28 | 0.00 | 4.66 | 0.00 | 8.61 | 0.00 |
|  | IFNL3 | 0.35 | 0.00 | 0.58 | 0.44 | 2.38 | 0.31 | 0.88 | 0.00 | 3.02 | 0.00 | 8.46 | 0.00 |
| ISGs | IFIT1 | 388.42 | 358.77 | 370.80 | 487.33 | 447.59 | 590.32 | 425.31 | 498.05 | 463.17 | 408.65 | 2790.57 | 367.50 |
|  | IRF7 | 278.50 | 283.73 | 320.43 | 284.00 | 339.99 | 383.89 | 399.07 | 363.67 | 399.93 | 432.29 | 966.54 | 305.31 |
|  | OAS2 | 172.67 | 236.24 | 178.18 | 222.85 | 287.61 | 208.20 | 252.85 | 296.10 | 292.36 | 378.90 | 2979.22 | 303.60 |
|  | MX1 | 588.48 | 620.75 | 624.79 | 647.52 | 758.95 | 800.13 | 839.47 | 867.29 | 728.29 | 811.68 | 3922.41 | 546.94 |
|  | RSAD2 | 204.76 | 216.53 | 228.73 | 272.67 | 313.84 | 348.31 | 365.12 | 393.68 | 274.53 | 269.56 | 948.75 | 210.54 |
|  | SLC44A4 | 1247.82 | 1171.72 | 1218.77 | 1046.17 | 1138.09 | 1128.19 | 1129.60 | 1106.06 | 1010.30 | 1142.19 | 1032.09 | 1298.09 |
|  | IFIH1 | 1052.81 | 1100.39 | 1134.78 | 1163.76 | 1235.36 | 1164.31 | 1223.66 | 1371.55 | 1189.70 | 1191.00 | 2492.69 | 1087.88 |
|  | GBP1 | 506.79 | 512.73 | 503.57 | 608.29 | 496.74 | 485.28 | 458.14 | 509.15 | 530.04 | 509.53 | 1151.35 | 488.92 |
|  | IFI44 | 689.16 | 741.40 | 789.19 | 803.61 | 963.68 | 1113.99 | 997.06 | 1052.67 | 785.42 | 782.39 | 1889.54 | 671.51 |
|  | IFI27 | 311.49 | 318.74 | 302.63 | 399.59 | 343.37 | 472.30 | 328.28 | 361.48 | 333.63 | 351.85 | 921.55 | 342.54 |
|  | IFI6 | 592.82 | 612.04 | 599.90 | 697.80 | 673.06 | 1010.20 | 692.26 | 752.25 | 729.19 | 775.17 | 2066.30 | 709.85 |
|  | ISG15 | 430.95 | 447.57 | 443.60 | 533.02 | 465.88 | 704.43 | 490.49 | 554.07 | 473.88 | 502.97 | 1260.48 | 435.91 |
|  | IFIT2 | 657.23 | 698.02 | 676.46 | 795.49 | 645.57 | 732.08 | 455.75 | 504.29 | 493.48 | 422.04 | 1465.16 | 413.27 |
|  | USP18 | 212.27 | 217.53 | 218.01 | 257.03 | 253.55 | 301.50 | 266.17 | 297.44 | 243.57 | 232.66 | 873.18 | 218.27 |
|  | IFIT3 | 648.15 | 656.89 | 747.61 | 858.17 | 810.13 | 1069.67 | 567.26 | 668.13 | 458.25 | 428.90 | 1900.07 | 420.64 |
|  | CMPK2 | 163.89 | 179.41 | 169.11 | 182.05 | 219.35 | 244.03 | 235.97 | 265.54 | 172.78 | 201.60 | 906.22 | 153.23 |
|  | XAF1 | 58.53 | 82.76 | 73.40 | 53.61 | 69.79 | 60.14 | 79.67 | 55.09 | 86.30 | 91.97 | 513.01 | 90.51 |
|  | IFITM1 | 27.68 | 34.25 | 21.94 | 27.89 | 28.53 | 53.49 | 26.88 | 34.91 | 34.59 | 35.75 | 182.01 | 34.33 |
|  | MX2 | 82.11 | 87.24 | 69.22 | 81.96 | 100.75 | 83.43 | 87.84 | 87.48 | 108.05 | 78.88 | 547.98 | 64.92 |

**Table S3.**
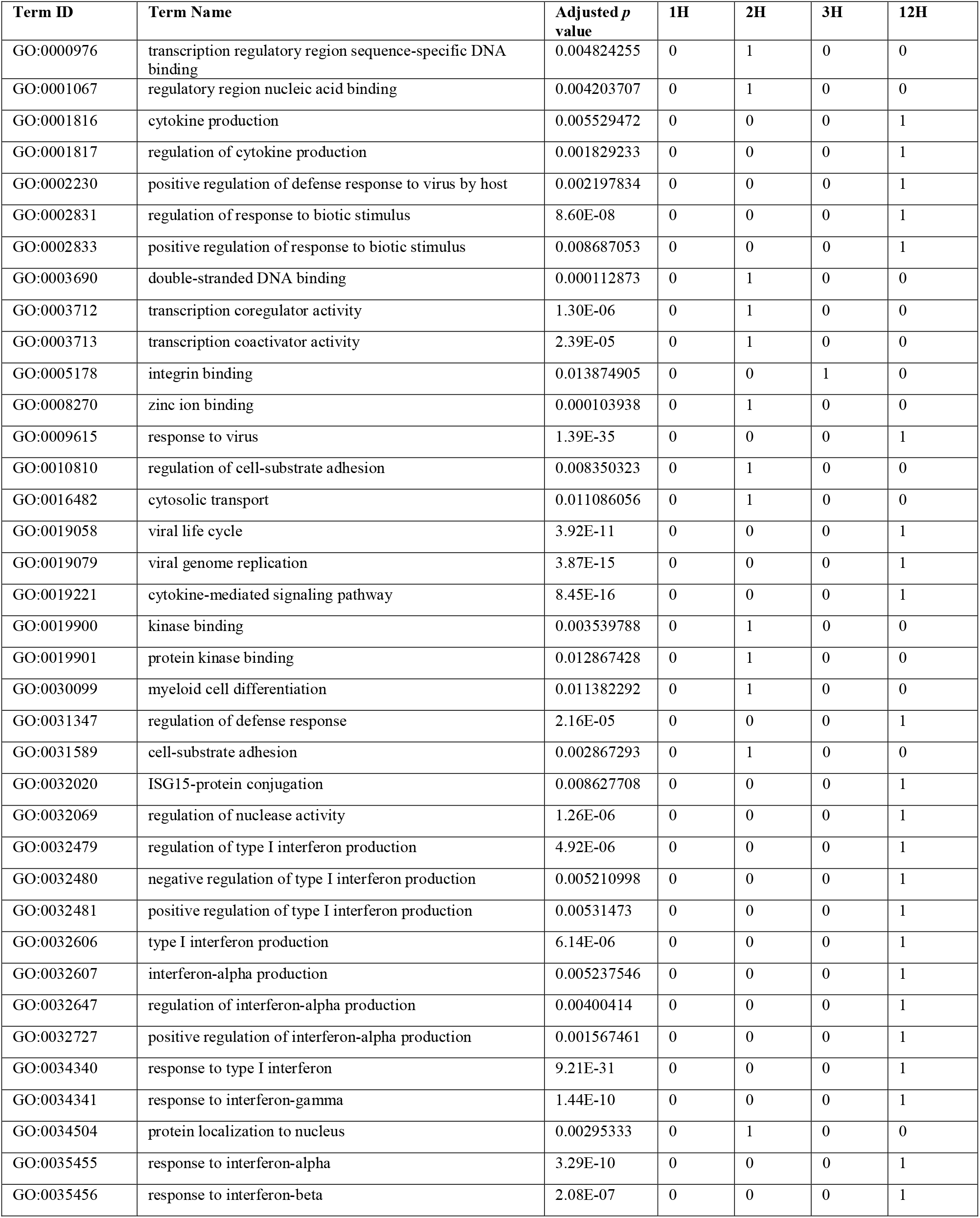
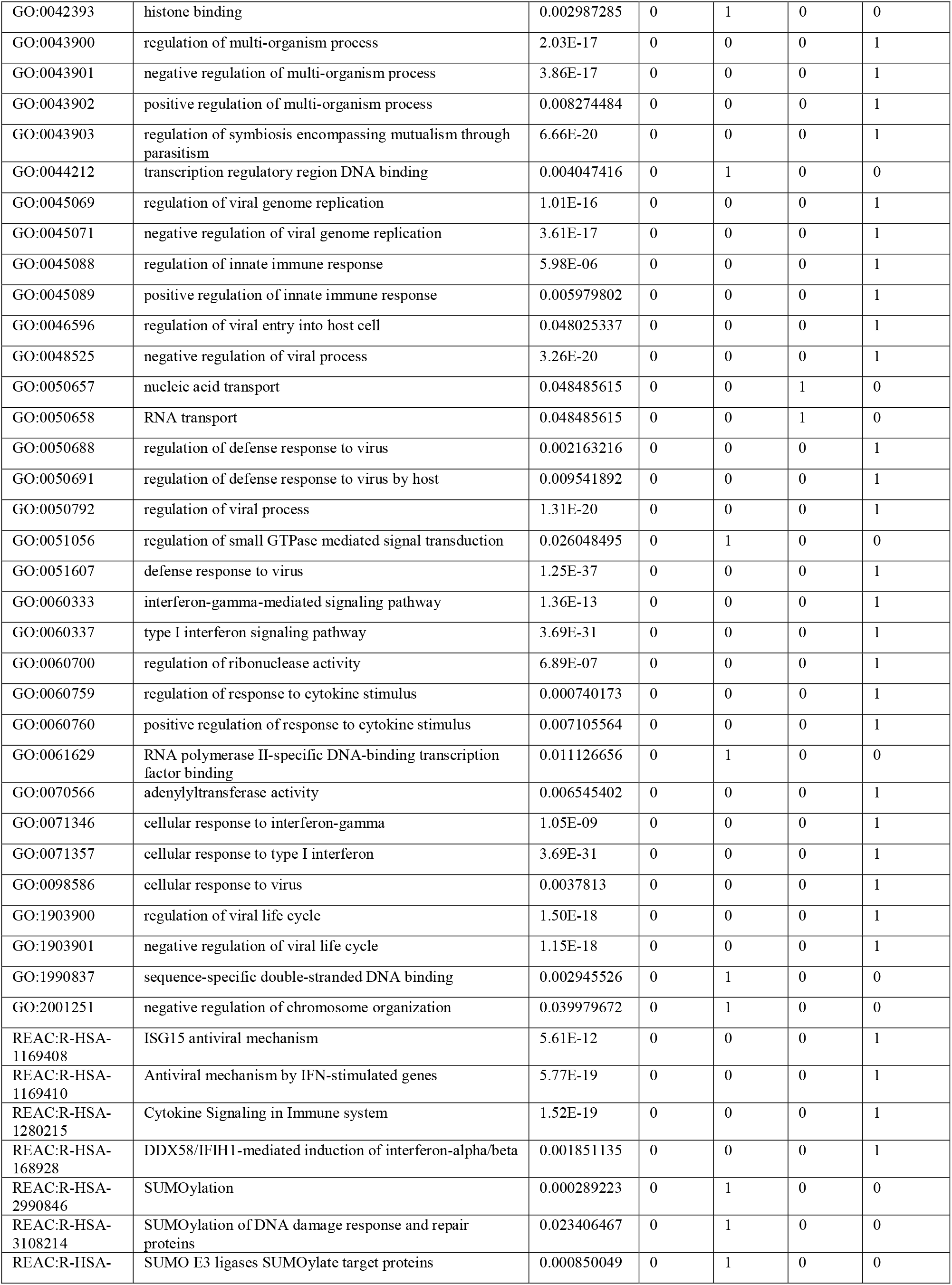
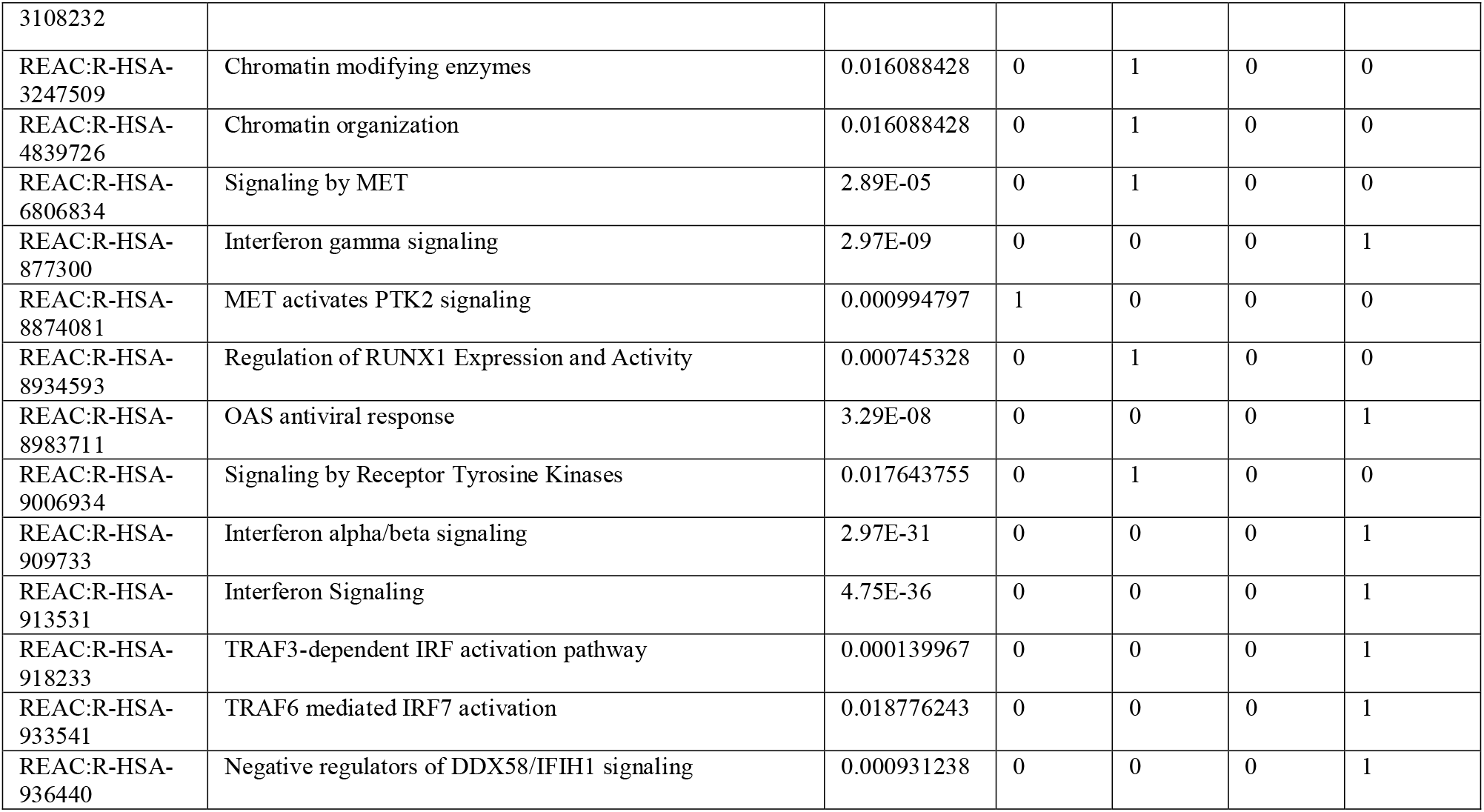
Pathway enrichment analysis. Significance was determined after FDR correction. H, hours post incubation; 0, non-significant; 1, significant. Related to Figure 1.

**Table S4.** COVID-19 patient serum sample history and sera from healthy controls, Related to Figure 2.

| Sample ID | Patient ID | Sample type | ICU admission | Outcome | Severity |
| --- | --- | --- | --- | --- | --- |
| CVD00845 | V097 | serum - acute | No | Discharged | Moderate |
| CVD01018 | V1282 | serum - acute | No | Discharged | Moderate |
| CVD01286 | V1718 | serum - acute | No | Discharged | Moderate |
| CVD01314 | V1716 | serum - acute | No | Discharged | Moderate |
| CVD01384 | V1826 | serum - acute | No | Discharged | Moderate |
| CVD01385 | V1804 | serum - acute | No | Discharged | Moderate |
| CVD01447 | V1830 | serum - acute | No | Discharged | Moderate |
| CVD01569 | V1939 | serum - acute | No | Discharged | Moderate |
| CVD01609 | V1999 | serum - acute | No | Discharged | Moderate |
| CVD01649 | V2000 | serum - acute | No | Discharged | Moderate |
| CVD00764 | V0861 | serum - acute | Yes | Transfer to rehab | Severe |
| CVD00768 | V0980 | serum - acute | Yes | Died | Severe |
| CVD00880 | V1100 | serum - acute | Yes | Discharged | Severe |
| CVD00907 | V1094 | serum - acute | Yes | Discharged | Severe |
| CVD01022 | V1293 | serum - acute | Yes | Died | Severe |
| CVD01339 | V1676 | serum - acute | Yes | Died | Severe |
| CVD01391 | V1714 | serum - acute | Yes | Discharged | Severe |
| CVD01433 | V1802 | serum - acute | Yes | Died | Severe |
| CVD01715 | V2071 | serum - acute | Yes | Died | Severe |
| CVD00864 | V0092 | serum - acute | Yes | Died | Severe |
| OM1 | NA | NA | NA | NA | Healthy |
| OM8035 | NA | NA | NA | NA | Healthy |
| OM908 | NA | NA | NA | NA | Healthy |
| OM920 | NA | NA | NA | NA | Healthy |
| OM921 | NA | NA | NA | NA | Healthy |

**Table S5.** Cytokine levels in healthy individuals and COVID-19 patient serum samples, Related to Figure 2. Data can be found in the Excel file labeled ‘Table S5’.

**Figure S1.**
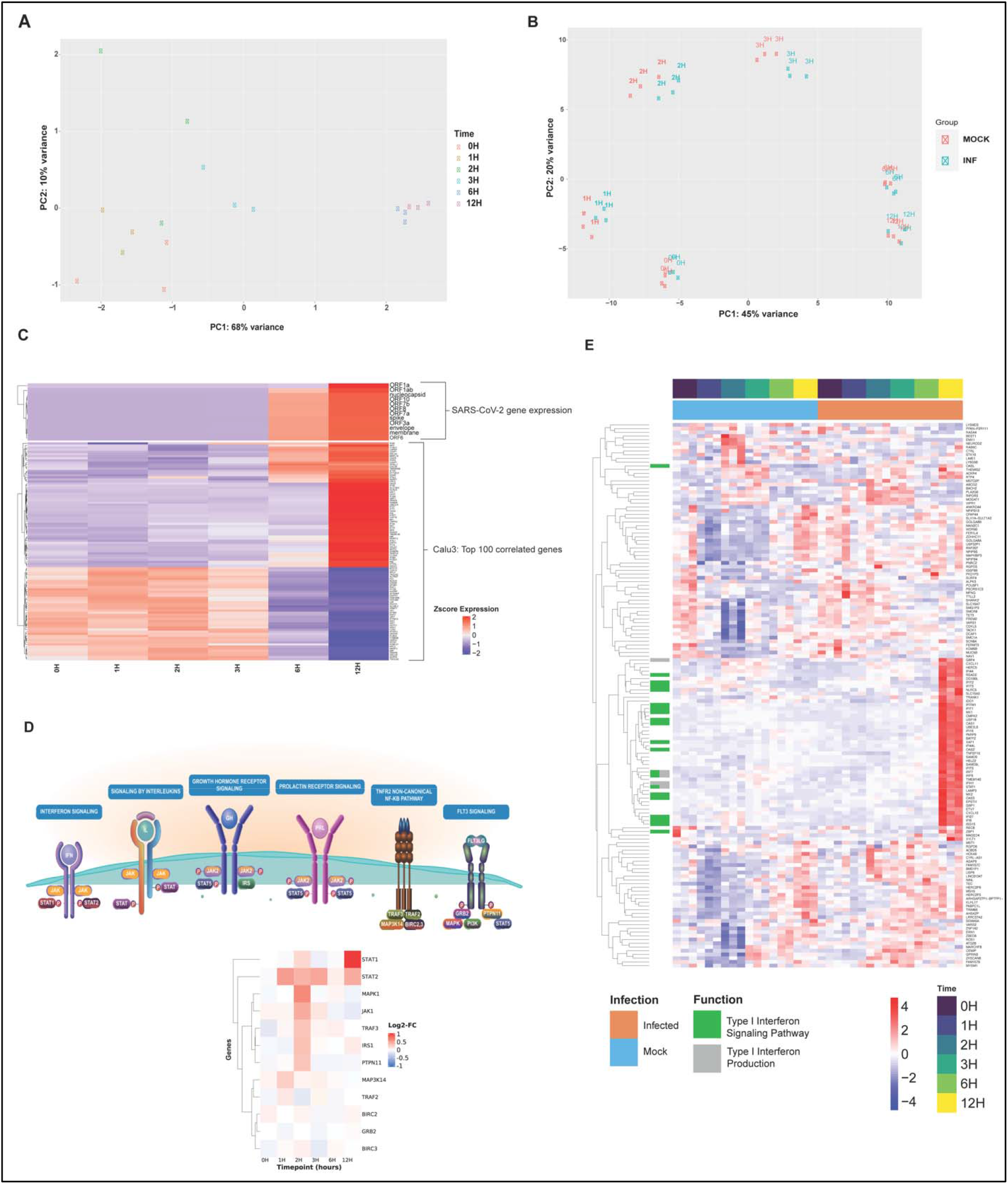
Transcript clustering and gene expression in Calu-3 cells infected with SARS-CoV-2, Related to Figure 1. To determine SARS-CoV-2 replication kinetics in human cells using RNA-seq, we infected human lung epithelial cells (Calu-3) at a multiplicity of infection (MOI) of 2. One hour post incubation, virus inoculum was replaced with cell growth media and the clock was set to zero hours. poly-A enriched RNA was extracted and sequenced at 0-, 1-, 2-, 3-, 6- and 12-hours post incubation (hpi). (**A**) SARS-CoV-2 genome, sub-genomic RNA and transcripts were detected in infected samples. PCA clustering was performed on quantified SARS-CoV-2 transcript levels in infected samples across time-points. Axes labels indicate the proportion of between-samples variance explained by the first two principal components. (**B**) PCA clustering was performed on quantified and filtered host gene transcripts in both SARS-CoV-2 infected (blue) and mock infected (red) samples across time-points (indicated in text for each data-point). Axes labels indicate the proportion of between-samples variance explained by the first two principal components. (**C**) Host gene expression that correlated with one or more viral transcripts over the course of infection are shown as z-score normalized expression (bottom), along with viral transcripts (top). Top 100 strongly correlated genes are represented here. (**D**) **Top**-Pathway schematic of REACTOME cytokine signalling pathway involving interferon alpha/beta/gamma signalling, and antiviral response mediated by interferon stimulated genes. **Bottom**-Heatmap of genes within REACTOME cytokine signalling pathway and their log_2_ transformed fold-change (FC) between SARS-CoV-2 infected and mock infected samples across all timepoints (0, 1, 2, 3, 6, 12 hours). (**E**) Larger version of Figure 1C. Cellular genes (n = 124) that are significantly up or downregulated (FDR-adjusted *p*<0.05; |log_2_FC| > 1) in SARS-CoV-2 infected cells, relative to mock infected cells at different times post incubation. Transcript levels are shown as z-score normalized expression (scaled by gene). H, hours post incubation. Mock, mock infected; INF, SARS-CoV-2 infected.

**Figure S2.**
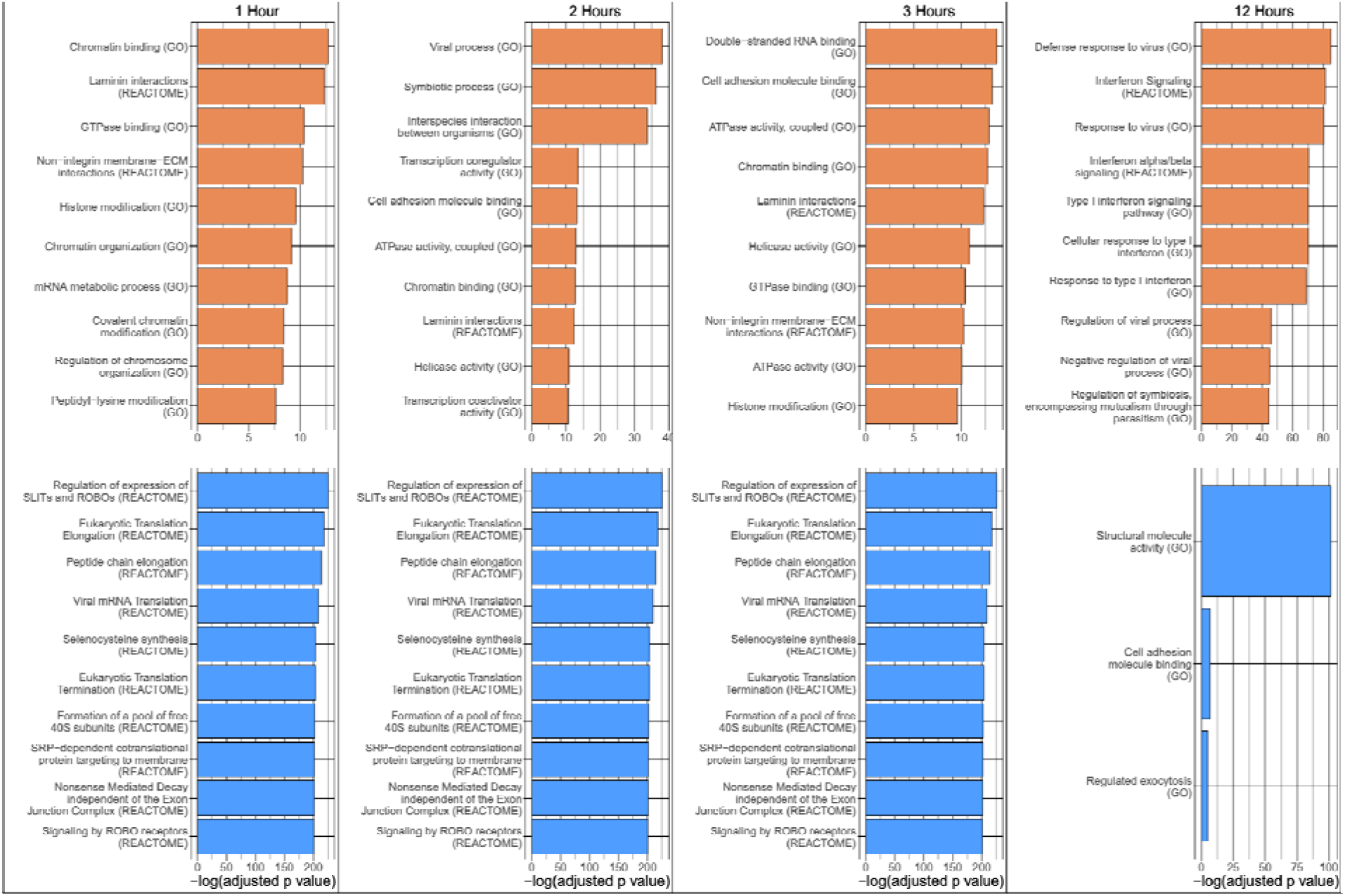
Top functional enrichments over time, Related to Figure 1. Top significantly (adjusted *p*<0.05) enriched ActivePathway GO terms and REACTOME enrichments for infection vs. mock at 1, 2, 3 and 12 hours post infection with SARS-CoV-2. Orange bars represent enriched terms associated with genes upregulated in infection vs. mock. Blue bars represent enriched terms associated with genes downregulated in infection vs. mock. 0 and 6 hour time points were omitted due to lack of sufficient numbers of differentially expressed genes.

**Figure S3.**
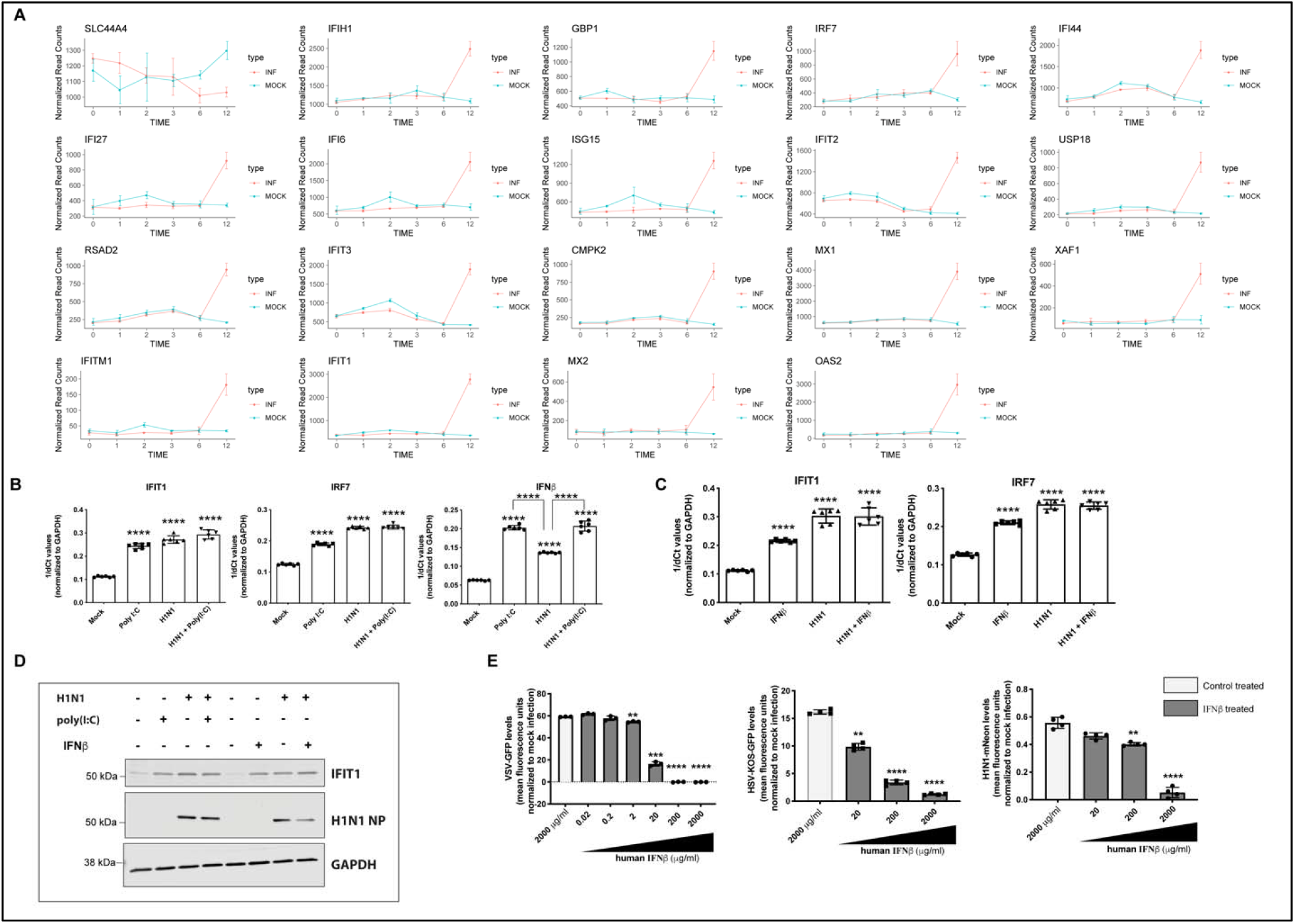
Infection-responsive gene expression profiles for ISGs, H1N1 infections in Calu-3 cells and characterization of recombinant human IFNβ1, Related to Figures 2 and 3. (**A**) ISG expression in Calu-3 cells with significantly different levels of transcript expression between mock (blue) and SARS-CoV-2 infected (red) samples at 12 hpi are shown (n = 3/time point). Normalized read counts per gene, across six time-points are represented here. Time indicated is in hours. Mock, mock infected; INF, SARS-CoV-2 infected. (**B**) *IFNβ*, *IFIT1* and *IRF7* transcript levels in Calu-3 cells that were infected with H1N1 (MOI = 1) or mock infected for 12 hours. Twelve hpi, cells were either transfected with 100 ng of poly(I:C) or mock transfected for 6 hours. *IFNβ*, *IFIT1* and *IRF7* transcript levels were normalized to *GAPDH* transcript levels (n = 6). (**C**) *IFIT1* and *IRF7* transcript levels in Calu-3 cells that were infected with H1N1 (MOI = 1) or mock infected for 12 hours. Twelve hpi, cells were either treated with 2 mg/ml recombinant IFNβ or mock treated for 6 hours. *IFIT1* and *IRF7* transcript levels were normalized to *GAPDH* transcript levels (n = 6). (**D**) H1N1 NP, IFIT1 and GAPDH protein expression in Calu-3 cells that were infected with H1N1 (MOI = 1) or mock infected for 12 hours. Twelve hpi, cells were either transfected or treated with poly (I:C) or recombinant IFNβ, respectively or mock transfected or treated for 6 hours (n=3). (**E**) Human fibroblast (THF) cells were treated with increasing concentrations of recombinant human IFNβ1 or mock treated with GFP containing media (control) for 6 hours. Cells were then infected with vesicular stomatitis virus (VSV-GFP), herpes simplex virus (HSV-KOS-GFP) or H1N1 influenza virus (H1N1-mNeon). VSV and HSV were engineered to express green fluorescent protein (GFP). H1N1 expressed mNeon that is detectable in the same wavelength as GFP. Nineteen hours post incubation, GFP or mNeon levels were measured in mock infected and virus infected cells as a surrogate for virus replication. VSV-GFP (n = 3), HSV-KOS-GFP (n = 4) and H1N1-mNeon (n = 4) replication in THF cells treated with IFNβ1 or mock treated with control, normalized to mock infection is shown above. Data are represented as mean ± SD, n=3, 4 or 6, *p*\*\*<0.01, ***<0.001 and ****<0.0001 (Student’s t test). GFP and mNeon expression is represented after normalization with mock infected cells. NP, nucleoprotein; hpi, hours post incubation.

**Figure S4.**
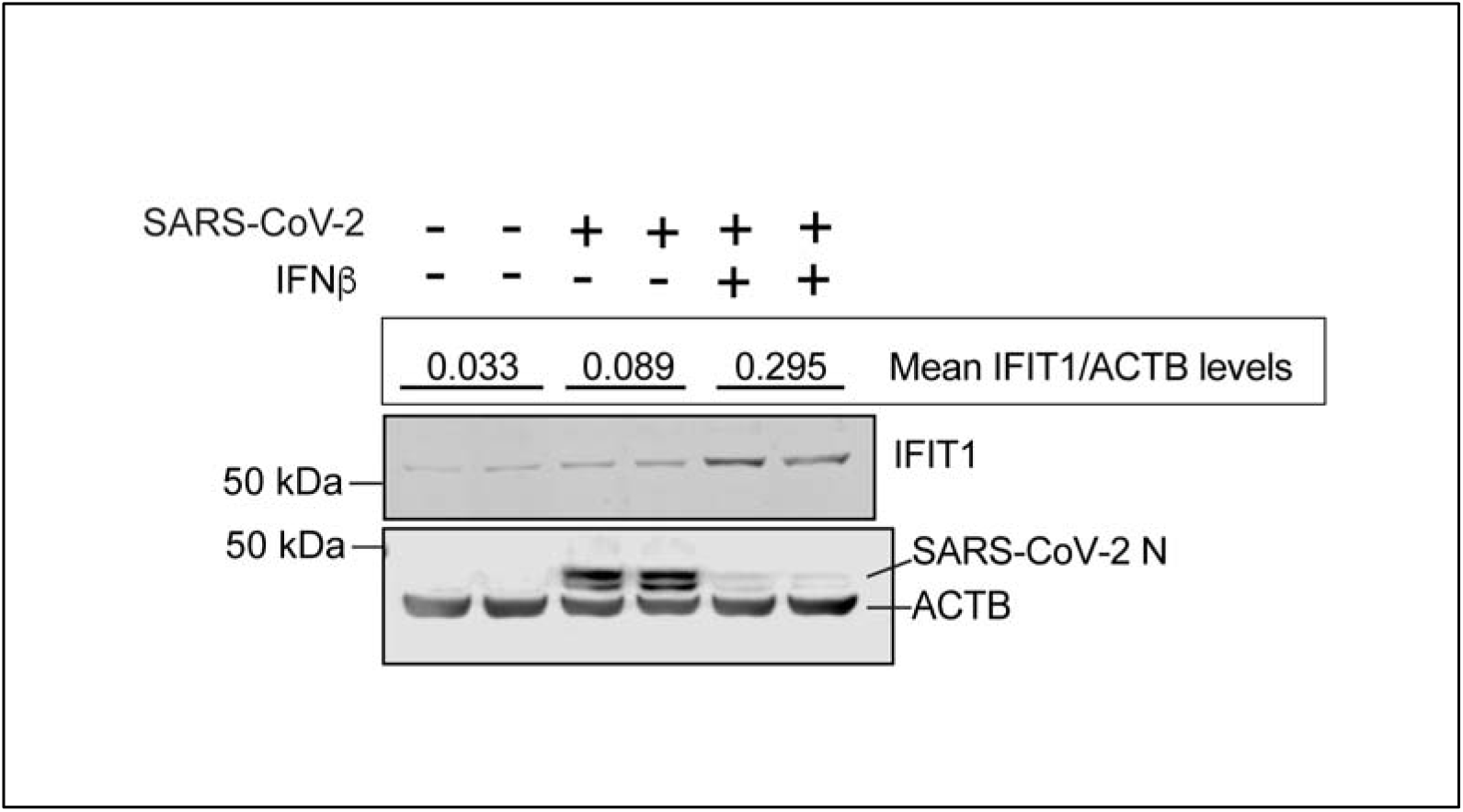
Protein expression in infected or treated Calu-3 cells, Related to Figure 4. IFIT1, SARS-CoV-2 N and ACTB protein expression in Calu-3 cells that were infected with SARS-CoV-2 or mock infected for 1 hr, followed by control or IFNβ treatment for 72 hours (n = 2). Data are represented as mean ± SD, n = 2. IFIT1 protein expression levels are expressed as ratios of IFIT1/ACTB levels. Blots were quantified using Image Studio (Li-COR) (n = 2).

## Notes

### Summary of Updates

In vitro immunoblot data and serum cytokine protein levels (Figures 2-4).

## References

Acharya, D., Liu, G., and Gack, M.U. (2020). Dysregulation of type I interferon responses in COVID-19. Nat Rev Immunol 20, 397–398.

Aguiar, J.A., Huff, R.D., Tse, W., Stampfli, M.R., McConkey, B.J., Doxey, A.C., and Hirota, J.A. (2019). Transcriptomic and barrier responses of human airway epithelial cells exposed to cannabis smoke. Physiol Rep 7, e14249.

Andrae, J., Gallini, R., and Betsholtz, C. (2008). Role of platelet-derived growth factors in physiology and medicine. Genes Dev 22, 1276–1312.

Arabi, Y.M., Asiri, A.Y., Assiri, A.M., Aziz Jokhdar, H.A., Alothman, A., Balkhy, H.H., AlJohani, S., Al Harbi, S., Kojan, S., Al Jeraisy, M., et al. (2020). Treatment of Middle East respiratory syndrome with a combination of lopinavir/ritonavir and interferon-beta1b (MIRACLE trial): statistical analysis plan for a recursive two-stage group sequential randomized controlled trial. Trials 21, 8.

Banerjee, A., Baid, K., and Mossman, K. (2019). Molecular Pathogenesis of Middle East Respiratory Syndrome (MERS) Coronavirus. Curr Clin Microbiol Rep 6, 139–147.

Banerjee, A., Nasir, J.A., Budylowski, P., Yip, L., Aftanas, P., Christie, N., Ghalami, A., Baid, K., Raphenya, A.R., Hirota, J.A., et al. (2020a). Isolation, Sequence, Infectivity, and Replication Kinetics of Severe Acute Respiratory Syndrome Coronavirus 2. Emerg Infect Dis 26.

Banerjee, A., Rapin, N., Bollinger, T., and Misra, V. (2017). Lack of inflammatory gene expression in bats: a unique role for a transcription repressor. Sci Rep 7, 2232.

Banerjee, A., Zhang, X., Yip, A., Schulz, K.S., Irving, A.T., Bowdish, D., Golding, B., Wang, L.F., and Mossman, K. (2020b). Positive Selection of a Serine Residue in Bat IRF3 Confers Enhanced Antiviral Protection. iScience 23, 100958.

Benjamini, Y., and Hochberg, Y. (1995). Controlling the False Discovery Rate: A Practical and Powerful Approach to Multiple Testing. Journal of the Royal Statistical Society: Series B (Methodological) 57, 289–300.

Blanco-Melo, D., Nilsson-Payant, B.E., Liu, W.C., Uhl, S., Hoagland, D., Moller, R., Jordan, T.X., Oishi, K., Panis, M., Sachs, D., et al. (2020). Imbalanced Host Response to SARS-CoV-2 Drives Development of COVID-19. Cell.

Chen, W., Srinath, H., Lam, S.S., Schiffer, C.A., Royer, W.E., Jr., and Lin, K. (2008). Contribution of Ser386 and Ser396 to activation of interferon regulatory factor 3. J Mol Biol 379, 251–260.

Chen, X., Yang, X., Zheng, Y., Yang, Y., Xing, Y., and Chen, Z. (2014). SARS coronavirus papain-like protease inhibits the type I interferon signaling pathway through interaction with the STING-TRAF3-TBK1 complex. Protein Cell 5, 369–381.

Couper, K.N., Blount, D.G., and Riley, E.M. (2008). IL-10: the master regulator of immunity to infection. J Immunol 180, 5771–5777.

de Wit, E., van Doremalen, N., Falzarano, D., and Munster, V.J. (2016). SARS and MERS: recent insights into emerging coronaviruses. Nat Rev Microbiol 14, 523–534.

Ding, Z., Fang, L., Jing, H., Zeng, S., Wang, D., Liu, L., Zhang, H., Luo, R., Chen, H., and Xiao, S. (2014). Porcine epidemic diarrhea virus nucleocapsid protein antagonizes beta interferon production by sequestering the interaction between IRF3 and TBK1. J Virol 88, 8936–8945.

Dong, E., Du, H., and Gardner, L. (2020). An interactive web-based dashboard to track COVID-19 in real time. Lancet Infect Dis 20, 533–534.

Fehr, A.R., and Perlman, S. (2015). Coronaviruses: an overview of their replication and pathogenesis. Methods Mol Biol 1282, 1–23.

Gordon, D.E., Jang, G.M., Bouhaddou, M., Xu, J., Obernier, K., White, K.M., O’Meara, M.J., Rezelj, V.V., Guo, J.Z., Swaney, D.L., et al. (2020). A SARS-CoV-2 protein interaction map reveals targets for drug repurposing. Nature 583, 459–468.

Grimminger, F., and Schermuly, R.T. (2010). PDGF receptor and its antagonists: role in treatment of PAH. Adv Exp Med Biol 661, 435–446.

Gu, Z., Eils, R., and Schlesner, M. (2016). Complex heatmaps reveal patterns and correlations in multidimensional genomic data. Bioinformatics 32, 2847–2849.

Hadjadj, J., Yatim, N., Barnabei, L., Corneau, A., Boussier, J., Pere, H., Charbit, B., Bondet, V., Chenevier-Gobeaux, C., Breillat, P., et al. (2020a). Impaired type I interferon activity and exacerbated inflammatory responses in severe Covid-19 patients. medRxiv.

Hadjadj, J., Yatim, N., Barnabei, L., Corneau, A., Boussier, J., Smith, N., Pere, H., Charbit, B., Bondet, V., Chenevier-Gobeaux, C., et al. (2020b). Impaired type I interferon activity and inflammatory responses in severe COVID-19 patients. Science 369, 718–724.

Hung, I.F.-N., Lung, K.-C., Tso, E.Y.-K., Liu, R., Chung, T.W.-H., Chu, M.-Y., Ng, Y.-Y., Lo, J., Chan, J., Tam, A.R., et al. (2020). Triple combination of interferon beta-1b, lopinavir#x2013; ritonavir, and ribavirin in the treatment of patients admitted to hospital with COVID-19: an open-label, randomised, phase 2 trial. The Lancet 395, 1695–1704.

Hunt, S.E., McLaren, W., Gil, L., Thormann, A., Schuilenburg, H., Sheppard, D., Parton, A., Armean, I.M., Trevanion, S.J., Flicek, P., et al. (2018). Ensembl variation resources. Database (Oxford) 2018.

Janeway, C.A., Jr., and Medzhitov, R. (2002). Innate immune recognition. Annu Rev Immunol 20, 197–216.

Jassal, B., Matthews, L., Viteri, G., Gong, C., Lorente, P., Fabregat, A., Sidiropoulos, K., Cook, J., Gillespie, M., Haw, R., et al. (2020). The reactome pathway knowledgebase. Nucleic Acids Res 48, D498–D503.

Jewell, N.A., Vaghefi, N., Mertz, S.E., Akter, P., Peebles, R.S., Jr., Bakaletz, L.O., Durbin, R.K., Flano, E., and Durbin, J.E. (2007). Differential type I interferon induction by respiratory syncytial virus and influenza a virus in vivo. J Virol 81, 9790–9800.

Jia, D., Rahbar, R., Chan, R.W., Lee, S.M., Chan, M.C., Wang, B.X., Baker, D.P., Sun, B., Peiris, J.S., Nicholls, J.M., et al. (2010). Influenza virus non-structural protein 1 (NS1) disrupts interferon signaling. PLoS One 5, e13927.

Jiang, H.W., Zhang, H.N., Meng, Q.F., Xie, J., Li, Y., Chen, H., Zheng, Y.X., Wang, X.N., Qi, H., Zhang, J., et al. (2020). SARS-CoV-2 Orf9b suppresses type I interferon responses by targeting TOM70. Cell Mol Immunol.

Katze, M.G., He, Y., and Gale, M., Jr. (2002). Viruses and interferon: a fight for supremacy. Nat Rev Immunol 2, 675–687.

Kawai, T., and Akira, S. (2006). Innate immune recognition of viral infection. Nat Immunol 7, 131–137.

Kochs, G., Garcia-Sastre, A., and Martinez-Sobrido, L. (2007). Multiple anti-interferon actions of the influenza A virus NS1 protein. J Virol 81, 7011–7021.

Koyama, S., Ishii, K.J., Coban, C., and Akira, S. (2008). Innate immune response to viral infection. Cytokine 43, 336–341.

Krueger, F. (2019). Trim Galore.

Lai, M.M. (1990). Coronavirus: organization, replication and expression of genome. Annu Rev Microbiol 44, 303–333.

Larabi, A., Devos, J.M., Ng, S.L., Nanao, M.H., Round, A., Maniatis, T., and Panne, D. (2013). Crystal structure and mechanism of activation of TANK-binding kinase 1. Cell Rep 3, 734–746.

Lei, X., Dong, X., Ma, R., Wang, W., Xiao, X., Tian, Z., Wang, C., Wang, Y., Li, L., Ren, L., et al. (2020). Activation and evasion of type I interferon responses by SARS-CoV-2. Nat Commun 11, 3810.

Li, H. (2013). Aligning sequence reads, clone.sequences. and assembly contigs with BWA-MEM. arXiv:13033997 [q-bioGN].

Li, H., and Durbin, R. (2009). Fast and accurate short read alignment with Burrows-Wheeler transform. Bioinformatics 25, 1754–1760.

Li, H., Handsaker, B., Wysoker, A., Fennell, T., Ruan, J., Homer, N., Marth, G., Abecasis, G., Durbin, R., and Genome Project Data Processing, S. (2009). The Sequence Alignment/Map format and SAMtools. Bioinformatics 25, 2078–2079.

Long, Q.X., Tang, X.J., Shi, Q.L., Li, Q., Deng, H.J., Yuan, J., Hu, J.L., Xu, W., Zhang, Y., Lv, F.J., et al. (2020). Clinical and immunological assessment of asymptomatic SARS-CoV-2 infections. Nat Med 26, 1200–1204.

Love, M.I., Huber, W., and Anders, S. (2014). Moderated estimation of fold change and dispersion for RNA-seq data with DESeq2. Genome Biol 15, 550.

Lu, X., Pan, J., Tao, J., and Guo, D. (2011). SARS-CoV nucleocapsid protein antagonizes IFN-beta response by targeting initial step of IFN-beta induction pathway, and its C-terminal region is critical for the antagonism. Virus Genes 42, 37–45.

Lucas, C., Wong, P., Klein, J., Castro, T.B.R., Silva, J., Sundaram, M., Ellingson, M.K., Mao, T., Oh, J.E., Israelow, B., et al. (2020). Longitudinal analyses reveal immunological misfiring in severe COVID-19. Nature 584, 463–469.

Lui, P.Y., Wong, L.Y., Fung, C.L., Siu, K.L., Yeung, M.L., Yuen, K.S., Chan, C.P., Woo, P.C., Yuen, K.Y., and Jin, D.Y. (2016). Middle East respiratory syndrome coronavirus M protein suppresses type I interferon expression through the inhibition of TBK1-dependent phosphorylation of IRF3. Emerg Microbes Infect 5, e39.

Mandel, M., Harari, G., Gurevich, M., and Achiron, A. (2020). Cytokine prediction of mortality in COVID19 patients. Cytokine 134, 155190.

Merico, D., Isserlin, R., Stueker, O., Emili, A., and Bader, G.D. (2010). Enrichment map: a network-based method for gene-set enrichment visualization and interpretation. PLoS One 5, e13984.

Mesev, E.V., LeDesma, R.A., and Ploss, A. (2019). Decoding type I and III interferon signalling during viral infection. Nat Microbiol 4, 914–924.

Minaker, R.L., Mossman, K.L., and Smiley, J.R. (2005). Functional inaccessibility of quiescent herpes simplex virus genomes. Virol J 2, 85.

Nasir, J.A., Kozak, R.A., Aftanas, P., Raphenya, A.R., Smith, K.M., Maguire, F., Maan, H., Alruwaili, M., Banerjee, A., Mbareche, H., et al. (2020). A Comparison of Whole Genome Sequencing of SARS-CoV-2 Using Amplicon-Based Sequencing, Random Hexamers and Bait Capture. Viruses 12.

Niemeyer, D., Zillinger, T., Muth, D., Zielecki, F., Horvath, G., Suliman, T., Barchet, W., Weber, F., Drosten, C., and Muller, M.A. (2013). Middle East respiratory syndrome coronavirus accessory protein 4a is a type I interferon antagonist. J Virol 87, 12489–12495.

Noyce, R.S., Collins, S.E., and Mossman, K.L. (2009). Differential modification of interferon regulatory factor 3 following virus particle entry. J Virol 83, 4013–4022.

Noyce, R.S., Taylor, K., Ciechonska, M., Collins, S.E., Duncan, R., and Mossman, K.L. (2011). Membrane perturbation elicits an IRF3-dependent, interferon‐independent antiviral response. J Virol 85, 10926–10931.

Paczkowska, M., Barenboim, J., Sintupisut, N., Fox, N.S., Zhu, H., Abd-Rabbo, D., Mee, M.W., Boutros, P.C., Drivers, P., Functional Interpretation Working, G., et al. (2020). Integrative pathway enrichment analysis of multivariate omics data. Nat Commun 11, 735.

Park, A., and Iwasaki, A. (2020). Type I and Type III Interferons - Induction, Signaling, Evasion, and Application to Combat COVID-19. Cell Host Microbe 27, 870–878.

Patro, R., Duggal, G., Love, M.I., Irizarry, R.A., and Kingsford, C. (2017). Salmon provides fast and bias-aware quantification of transcript expression. Nat Methods 14, 417–419.

Perlman, S., and Netland, J. (2009). Coronaviruses post-SARS: update on replication and pathogenesis. Nat Rev Microbiol 7, 439–450.

Pilz, A., Ramsauer, K., Heidari, H., Leitges, M., Kovarik, P., and Decker, T. (2003). Phosphorylation of the Stat1 transactivating domain is required for the response to type I interferons. EMBO Rep 4, 368–373.

Pripp, A.H., and Stanisic, M. (2014). The correlation between pro- and anti-inflammatory cytokines in chronic subdural hematoma patients assessed with factor analysis. PLoS One 9, e90149.

RCoreTeam (2017). R: A language and environment for statistical computing.

Reimand, J., Isserlin, R., Voisin, V., Kucera, M., Tannus-Lopes, C., Rostamianfar, A., Wadi, L., Meyer, M., Wong, J., Xu, C., et al. (2019). Pathway enrichment analysis and visualization of omics data using g:Profiler, GSEA, Cytoscape and EnrichmentMap. Nat Protoc 14, 482–517.

Sadiq, M.A., Hanout, M., Sarwar, S., Hassan, M., Do, D.V., Nguyen, Q.D., and Sepah, Y.J. (2015). Platelet derived growth factor inhibitors: A potential therapeutic approach for ocular neovascularization. Saudi J Ophthalmol 29, 287–291.

Sawicki, S.G., Sawicki, D.L., and Siddell, S.G. (2007). A contemporary view of coronavirus transcription. J Virol 81, 20–29.

Schoggins, J.W. (2019). Interferon-Stimulated Genes: What Do They All Do? Annu Rev Virol 6, 567–584.

Schoggins, J.W., and Rice, C.M. (2011). Interferon-stimulated genes and their antiviral effector functions. Curr Opin Virol 1, 519–525.

Schulz, K.S., and Mossman, K.L. (2016). Viral Evasion Strategies in Type I IFN Signaling - A Summary of Recent Developments. Front Immunol 7, 498.

Shannon, P., Markiel, A., Ozier, O., Baliga, N.S., Wang, J.T., Ramage, D., Amin, N., Schwikowski, B., and Ideker, T. (2003). Cytoscape: a software environment for integrated models of biomolecular interaction networks. Genome Res 13, 2498–2504.

Shin, D., Mukherjee, R., Grewe, D., Bojkova, D., Baek, K., Bhattacharya, A., Schulz, L., Widera, M., Mehdipour, A.R., Tascher, G., et al. (2020). Papain-like protease regulates SARS-CoV-2 viral spread and innate immunity. Nature.

Siu, K.L., Yeung, M.L., Kok, K.H., Yuen, K.S., Kew, C., Lui, P.Y., Chan, C.P., Tse, H., Woo, P.C., Yuen, K.Y., et al. (2014). Middle east respiratory syndrome coronavirus 4a protein is a double-stranded RNA-binding protein that suppresses PACT-induced activation of RIG-I and MDA5 in the innate antiviral response. J Virol 88, 4866–4876.

Soneson, C., Love, M.I., and Robinson, M.D. (2015). Differential analyses for RNA-seq: transcript-level estimates improve gene-level inferences. F1000Research 4.

Steen, H.C., and Gamero, A.M. (2013). STAT2 phosphorylation and signaling. JAKSTAT 2, e25790.

The Gene Ontology, C. (2019). The Gene Ontology Resource: 20 years and still GOing strong. Nucleic Acids Res 47, D330–D338.

Trouillet-Assant, S., Viel, S., Gaymard, A., Pons, S., Richard, J.C., Perret, M., Villard, M., Brengel-Pesce, K., Lina, B., Mezidi, M., et al. (2020). Type I IFN immunoprofiling in COVID-19 patients. J Allergy Clin Immunol.

Wickham, H. (2009). ggplot2: Elegant graphics for data analysis (New York: Springer-Verlag).

Wilk, A.J., Rustagi, A., Zhao, N.Q., Roque, J., Martinez-Colon, G.J., McKechnie, J.L., Ivison, G.T., Ranganath, T., Vergara, R., Hollis, T., et al. (2020). A single-cell atlas of the peripheral immune response in patients with severe COVID-19. Nat Med 26, 1070–1076.

Xia, H., Cao, Z., Xie, X., Zhang, X., Yun-Chung Chen, J., Wang, H., Menachery, V.D., Rajsbaum, R., and Shi, P.-Y. (2020). Evasion of type-I interferon by SARS-CoV-2. Cell Reports.

Xing, Y., Chen, J., Tu, J., Zhang, B., Chen, X., Shi, H., Baker, S.C., Feng, L., and Chen, Z. (2013). The papain-like protease of porcine epidemic diarrhea virus negatively regulates type I interferon pathway by acting as a viral deubiquitinase. J Gen Virol 94, 1554–1567.

Xu, K., Cai, H., Shen, Y., Ni, Q., Chen, Y., Hu, S., Li, J., Wang, H., Yu, L., Huang, H., et al. (2020). Translation: Management of Coronavirus Disease 2019 (COVID-19): Experience in Zhejiang Province, China. Infectious Microbes and Diseases 2, 55–63.

Yamakawa, S., and Hayashida, K. (2019). Advances in surgical applications of growth factors for wound healing. Burns Trauma 7, 10.

Yang, Y., Zhang, L., Geng, H., Deng, Y., Huang, B., Guo, Y., Zhao, Z., and Tan, W. (2013). The structural and accessory proteins M,ORF 4a, ORF 4b and ORF 5 of Middle East respiratory syndrome coronavirus (MERS-CoV) are potent interferon antagonists. Protein Cell 4, 951–961.

Zhou, P., Yang, X.L., Wang, X.G., Hu, B., Zhang, L., Zhang, W., Si, H.R., Zhu, Y., Li, B., Huang, C.L., et al. (2020a). A pneumonia outbreak associated with a new coronavirus of probable bat origin. Nature 579, 270–273.

Zhou, Z., Ren, L., Zhang, L., Zhong, J., Xiao, Y., Jia, Z., Guo, L., Yang, J., Wang, C., Jiang, S., et al. (2020b). Heightened Innate Immune Responses in the Respiratory Tract of COVID-19 Patients. Cell Host Microbe 27, 883–890 e882.

